# An in situ generated CAR-M with IFN-γ and negative dominance Sirpα isoform augments hepatocellular carcinoma immunotherapy

**DOI:** 10.64898/2025.12.08.693087

**Authors:** Xiang Li, Jing Hu, Tian Zhang, Hongceng Li, Tao Wang, Yuxin Liu, Lan Zhang, Yonglong Wei, Shi Wei, Panpan Zhang

## Abstract

**Background and Aims:** Chimeric antigen receptor (CAR) T cells have shown strong efficacy in hematological cancers but limited success in solid tumors. Macrophages, with their natural ability to infiltrate tumors, modulate immunity, and phagocytose cancer cells, offer a promising alternative when engineered with CARs. While studies have demonstrated the feasibility and anti-tumor activity of CAR macrophages (CAR-M), enhancing their persistence and phagocytic capacity remains a key challenge.

**Methods and Results:** Building on first-generation CD3ζ-based CAR-M, we developed a novel CAR targeting human GPC3, incorporating IFN-γ and the extracellular domain of SIRPα (SIRPα^ECD^) to improve CAR-M persistence and block the CD47–SIRPα immune checkpoint. Following delivery via lipid nanoparticle-encapsulated mRNA (LNP-mRNA), the self-secreted IFN-γ sustained M1 polarization through phospho-STAT1 activation. Meanwhile, the ectopically expressed SIRPα^ECD^ competitively bound to CD47 on tumor cells, thereby blocking the endogenous SIRPα–SHP2 interaction in a dominant-negative manner. This design enhanced pro-inflammatory activity and anti-tumor efficacy compared to CD3ζ-only CAR-M. Single-cell RNA sequencing and cellular analysis showed that in situ programmed CAR-M reprogrammed the tumor microenvironment toward inflammation in a murine HCC model. Moreover, CAR-M derived from human peripheral blood mononuclear cells (PBMCs) effectively phagocytosed human HCC organoids while sparing healthy tissues, indicating clinical potential.

**Conclusions:** Collectively, our work presents a novel CAR design that enhances phagocytic function and sustains anti-tumor activity, offering a promising strategy for human solid tumor immunotherapy.

## Introduction

Traditional CAR-T cell therapies have demonstrated remarkable therapeutic efficacy in the treatment of B cell malignancies, with several approved CAR-T therapies targeting antigens such as CD19 and BCMA. However, the development of effective CAR-T therapies for solid tumors remains a significant challenge. This is largely due to the limited infiltration and functional activity of CAR-T cells within solid tumor tissues, as well as the immunosuppressive nature of the tumor microenvironment (TME) [1, 2]. As a key component of the innate immune system, macrophages are capable of penetrating the physical barriers of solid tumors, engulfing tumor cells, and modulating the immune microenvironment through antigen presentation to activate T cells [3–5]. Therefore, although still in its early stages, CAR-M therapy is increasingly being recognized as a promising alternative immunotherapeutic strategy with great potential for clinical application in the treatment of solid tumors.

HCC is the most common type of liver cancer and ranks as the fourth leading cause of cancer-related mortality worldwide [6]. The tumor microenvironment in HCC is notably infiltrated by macrophages, which are referred to as tumor-associated macrophages (TAMs) [7, 8]. In most cases, TAMs exhibit an M2-like polarization state and secrete various anti-inflammatory cytokines, such as IL-10 and TGF-β, contributing to the formation of an immunosuppressive TME [9, 10]. This immunosuppressive state often renders conventional immunotherapies against HCC ineffective [11]. Although numerous studies have focused on adoptive CAR-M therapies using macrophages generated in vitro or ex vivo, the labor-intensive and costly manufacturing processes limit their broad clinical application. Recently, an in vivo programming strategy for generating CAR-engineered immune cells has attracted increasing attention [12]. Given the high infiltration and abundance of tissue-resident macrophages in the liver, in vivo CAR-M therapy, which enables the reprogramming of endogenous macrophages in situ, is undoubtedly a promising immunotherapeutic approach for HCC.

The earliest CAR structure design of CAR-M consists of an antigen-binding region, a hinge and transmembrane domain, and the CD3ζ intracellular signaling domain [13]. A subsequent study proposed a CD3ζ-TIR dual-signaling CAR structure capable of continuously activating toll-like receptor 4 (TLR4) signaling, thereby helping macrophages maintain the M1 polarization state [14]. Although both generations of CAR-M exhibit strong anti-cancer capabilities, maintaining the persistence of CAR-M remains one of the major challenges [4, 15]. Compounding the issue, tumor cells frequently express CD47, which triggers the “don’t eat me” signal to inhibit macrophage phagocytosis [16, 17], further limiting the anti-tumor efficacy of CAR-M.

In this study, building upon the CD3ζ intracellular signaling domain of the first-generation CAR-M, we designed a novel CAR structure incorporating two additional components: IFN-γ and the extracellular domain of SIRPα (SIRPα^ECD^). These elements were introduced to prolong CAR-M persistence and inhibit the CD47-SIRPα signaling axis, respectively. Both in vitro and in vivo data demonstrated that our dual-functional IFN-γ-SIRPα^ECD^ CAR-M exhibited enhanced persistence and stronger anti-tumor activity compared to previous designs. Further cellular analysis and single-cell RNA-seq results indicated that our in situ programming CAR-M strategy could reshape the TME of HCC, promoting a sustained pro-inflammatory state. Notably, our CAR structure is also applicable to human macrophages, as CAR-M effectively and specifically eliminated human liver tumor organoids. These findings provide a proof-of-concept for the next-generation CAR-M strategy, which enhances persistence and anti-tumor activity by integrating multiple functional modules.

## Results

### A novel Car design structure that enables macrophage activation

As a specifically and highly expressed tumor antigen in hepatocellular carcinoma, GPC3 is correlated with the overall survival of HCC patients (Fig. S1A–C), and has been recognized as a promising target for various ongoing immunotherapies. The single-chain antibody fragment (scFv) targeting human GPC3 was derived from previous studies [18]. Inspired by prior findings, we introduced two novel components into the CAR structure to enhance the anti-tumor activity of CAR-M. One of the major challenges in CAR-M development is improving persistence to ensure a sustained anti-tumor effect [15]. Therefore, we first overexpressed mouse Ifn-γ in our CAR construct to induce a prolonged pro-inflammatory response in CAR-M (Fig. 1A). Second, the CD47–SIRPα axis is well established as a key mediator of the “don’t eat me” signal within the TME [19]. To counteract this inhibitory pathway in CAR-M, we incorporated the Sirpα^ECD^ domain to competitively bind Cd47, which is highly expressed on tumor cells (Fig. S1D, Fig. 1A), thereby attenuating the interaction between the immunoreceptor tyrosine-based inhibitory motif (ITIM) and SHP1/2 [19, 20]. To evaluate the anti-tumor efficacy of our novel CAR design, we selected the previously reported CD3ζ-only CAR-M structure as a control. The corresponding CAR mRNAs were encapsulated into LNPs, designated LNP1 and LNP2 respectively (Fig. 1A). Both IFN-γ and TLR4 signaling pathways are known to activate macrophages. According to previous studies, TLR4 agonists can induce IFN-γ expression, while IFN-γ has been shown to suppress TLR4-mediated transcription in macrophages [21, 22], suggesting the presence of a potential negative feedback loop. To investigate whether elevated IFN-γ secretion is more effective than TLR4 activation in CAR-M, we treated macrophages with lipopolysaccharide (LPS), a pathogen-associated molecular pattern (PAMP), to activate TLR4 signaling, both with and without LNP transfection.

**Figure 1.**
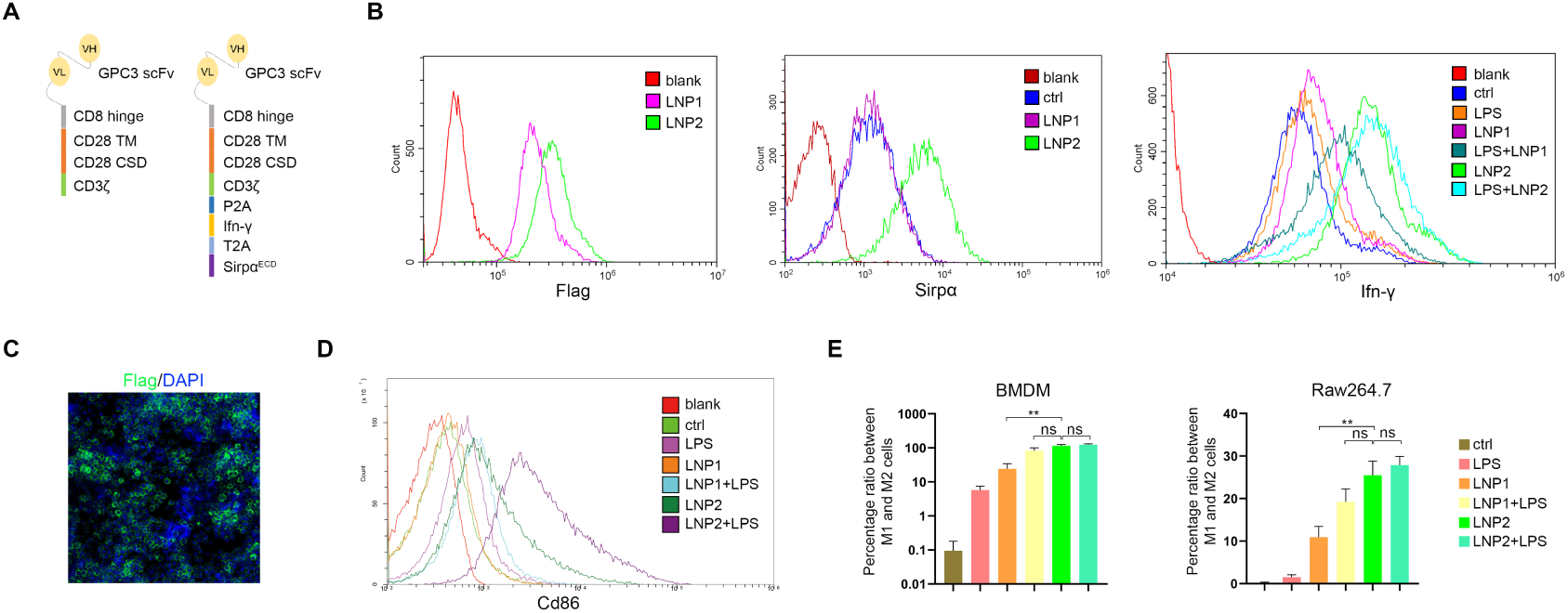
A novel CAR construct incorporating Ifn-γ and Sirpα^ECD^ functional modules demonstrates enhanced macrophage activation. (A) Schematic representation of two CAR constructs targeting the human GPC3 antigen. The constructs consist of a GPC3-specific scFv, a mouse Cd8 hinge region, a mouse Cd28 transmembrane (TM) domain and co-stimulatory domain (CSD), and a mouse Cd3ζ signaling domain. The mouse Ifn-γ coding sequence and Sirpα^ECD^ isoform were inserted into the CAR construct and separated by a 2A peptide to ensure their co-expression. (B) Flow cytometric analysis of signal intensity in BMDM cells following transfection with LNP1 or LNP2. Prior to transfection, BMDMs were polarized to an M1 phenotype. The blank group received no treatment, while the control (ctrl) group was transfected with empty LNP lacking mRNA. (C) Immunofluorescence imaging of CAR-M cells derived from BMDMs. A Flag tag was introduced downstream of the Cd3ζ domain to confirm CAR expression. (D) Flow cytometric assessment of Cd86 expression levels in BMDM-derived CAR-M cells transfected with LNP1 or LNP2. (E) Comparative and statistical analysis of flow cytometry data indicating the M1/M2 polarization ratio in BMDM and Raw264.7 cells after LNP or LPS treatment. \**p* <0.05, \*\**p* <0.01, ns, not significant.

We firstly transfected LNP-mRNA into the mouse macrophage cell line Raw264.7 and mouse bone marrow derived macrophage (BMDM). RT-qPCR, flow cytometry, and Western blot analyses confirmed high expression levels of the key CAR components in macrophages (Fig. S2A–C, Fig. 1B). Immunofluorescence staining demonstrated that the CAR protein was localized at the cell membrane (Fig. 1C). Interestingly, LPS treatment only moderately increased Ifn-γ expression in both the control and LNP1 groups (Fig. S2A, B, Fig. 1B). Next, we assessed the impact of CAR mRNA on macrophage polarization. The mRNA expression levels of M1-specific markers, including *iNos* and *Il6*, were significantly upregulated, among which LNP2 armed CAR-M displayed higher expression level than LPS treatment groups to some extent (Fig. S2A, B). Sequentially, BMDM cells arming by LNP1 or LNP2 were collected at a regular time interval. The time-course records of qRT-PCR results indicated that LNP1 derived CAR-M maintained the pro-inflammatory status for about 10 days, whereas LNP2 derived CAR-M could sustain this active phenotype for more than 16 days (Fig. S2D). Furthermore, flow cytometric analysis of the M1/M2 surface markers Cd86 and Cd206 revealed that both LNP1 and LNP2 promoted CD86 expression at the expense of CD206, with LNP2 showing significantly greater efficacy than LNP1 (Fig. S3A, B, Fig. 1D, E). Additionally, LPS treatment further enhanced M1 polarization in LNP1 CAR-M, but did not surpass the effect of LNP2 (Fig. S3A, B, Fig. 1D, E).

### CAR-M with the new CAR structure exhibited enhanced anti-tumor activity

Then, a mouse hepatocarcinoma cell line, Hepa1-6, was transfected with a lentiviral vector overexpressing human GPC3 and GFP (Fig. 2A). To further characterize the secretory profile of CAR-M, we co-cultured Hepa1-6-GPC3 tumor cells with BMDMs armed with either LNP1 or LNP2, and collected the culture supernatants for mass spectrometric analysis. The results revealed that the different treatment groups exhibited both shared and distinct secreted proteins involved in extracellular matrix remodeling, cell communication, and immune response pathways (Fig. S4A, B). Furthermore, we identified the most significantly enriched differentially expressed proteins (DEPs) in the LNP2-armed CAR-M group compared to the other groups, demonstrating that cytokines such as Ifn-γ, Il12, Il18, and Ccl8, as well as associated signaling pathways including the innate immune response and Jak-Stat signaling, were markedly activated upon LNP2 treatment (Fig. S4C, D, Fig. 2B).

**Figure 2.**
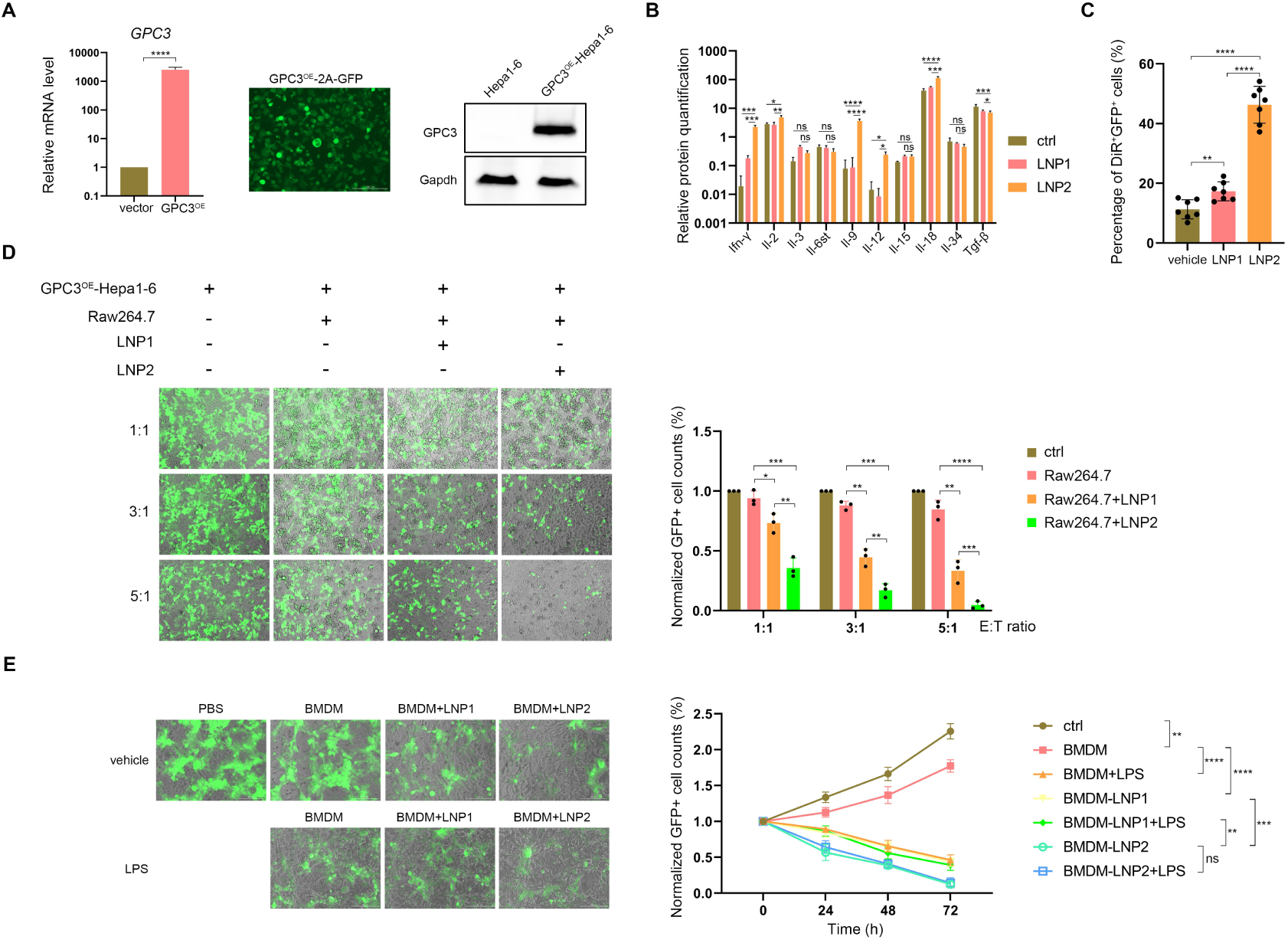
CAR-M cells effectively eliminate GPC3-overexpressing tumor cells in vitro. (A) qRT-PCR, immunofluorescence, and western blot analyses confirmed GPC3 mRNA and protein expression in Hepa1-6 cells. GPC3 overexpression (GPC3^OE^) was induced by lentiviral transduction with a 2A-GFP vector, with empty vector-transduced cells serving as the control. (B) Quantification of inflammatory cytokines secreted by BMDM-derived CAR-M cells transfected with LNP1 or LNP2, analyzed via mass spectrometry. (C) Comparative and statistical analysis of flow cytometry data assessing the phagocytic capacity of CAR-M cells. CAR-Ms were generated from BMDMs transfected with LNP1 or LNP2. Following labeling with the lipophilic dye DiR, CAR-Ms were co-cultured with GFP-expressing Hepa1-6 cells. Double-positive DiR and GFP cells were considered as macrophages that had engulfed tumor cells. (D) Live-cell imaging of GPC3^OE^-Hepa1-6 cells after 24 hours co-culturing with CAR-Ms at varying E:T ratios, along with quantification of GFP+ cells across treatment groups. CAR-Ms were derived from BMDMs, and GFP+ cell counts were determined using ImageJ software. (E) live-cell imaging of GPC3^OE^-Hepa1-6 cells after 72 hours co-culture with CAR-Ms at 1:1 E:T ratio, and time-course quantification of GFP+ cells across treatment groups. The left panel shows the co-culture status at 72 hours, while the right panel presents the time-dependent changes in GFP^+^ cell counts. The above statistical data were presented as mean ± s.e.m., and t-test analysis or two-way ANOVA analysis was used to calculate the significance. \**p* < 0.05; \*\**p* < 0.01; \*\*\**p* < 0.001; \*\*\*\**p* < 0.0001. ns, not significant.

Next, we compared the phagocytic activity of CAR-M with different design. After labeling with a lipophilic membrane fluorescent dye, DiR, CAR-M was co-cultured with Hepa1-6 tumor cells with or without GPC3 overexpression. After 6 hours of co-culture, CAR-M that had engulfed GFP^+^ tumor cells displayed double fluorescence positivity under ideal conditions. Flow cytometric results revealed that both LNP1-and LNP2-armed CAR-M specifically recognized the GPC3 antigen, while the LNP2 group exhibited significantly higher tumor cell engulfment (Fig. S5A, Fig. 2C).

We then evaluated the direct tumor-killing effect of CAR-M using live-cell imaging. LNP1- or LNP2-armed CAR-M was co-cultured with Hepa1-6 tumor cells at various effector-to-target (E:T) ratios ranging from 5:1 to 1:1, and GFP+ tumor cells were monitored at regular intervals. Both LNP1 and LNP2 demonstrated effective tumor-killing activity in a dose-dependent manner after 24 hours of co-culture (Fig. 2D). After 72 hours, LNP2-armed CAR-M, whether derived from Raw264.7 cells or BMDMs, exhibited significantly stronger anti-tumor activity compared to LNP1 or LNP1 combined with LPS, even at a 1:1 E:T ratio (Fig. S5B, Fig. 2E), suggesting that our novel CAR design enables macrophages to maintain a persistent activated state.

### Ifn-γ and Sirpα^ECD^ functions synergistically to potentiate anti-tumor activity of CAR-M

IFN-γ exerts pleiotropic anti-tumor effects within the TME, including initiating tumor cell apoptosis in conjunction with granzyme B and perforin, as well as inducing MHC class I (MHC-I) upregulation in tumor cells [23, 24]. Therefore, we collected the supernatant from CAR-M cultures and assessed its impact on tumor cell apoptosis and Mhc-I expression. The results showed that the supernatant from LNP2-armed CAR-M induced more pronounced tumor cell apoptosis and significantly upregulated Mhc-I expression (Fig. 3A, B), confirming the functional role of Ifn-γ. Moreover, Ifngr downstream Jak-Stat signaling was markedly activated upon LNP2 treatment (Fig. 3C). Meanwhile, analysis of the LPS-Tlr4 axis downstream Nf-κb signaling revealed that LNP2 did not interfere with LPS-mediated Nf-κb activation (Fig. 3C). These findings suggest that Ifn-γ–Jak-Stat and LPS–Tlr4–Nf-κb signaling pathways largely operate in parallel within macrophages, and that Ifn-γ–mediated macrophage activation contributes to a pleiotropic anti-tumor response.

**Figure 3.**
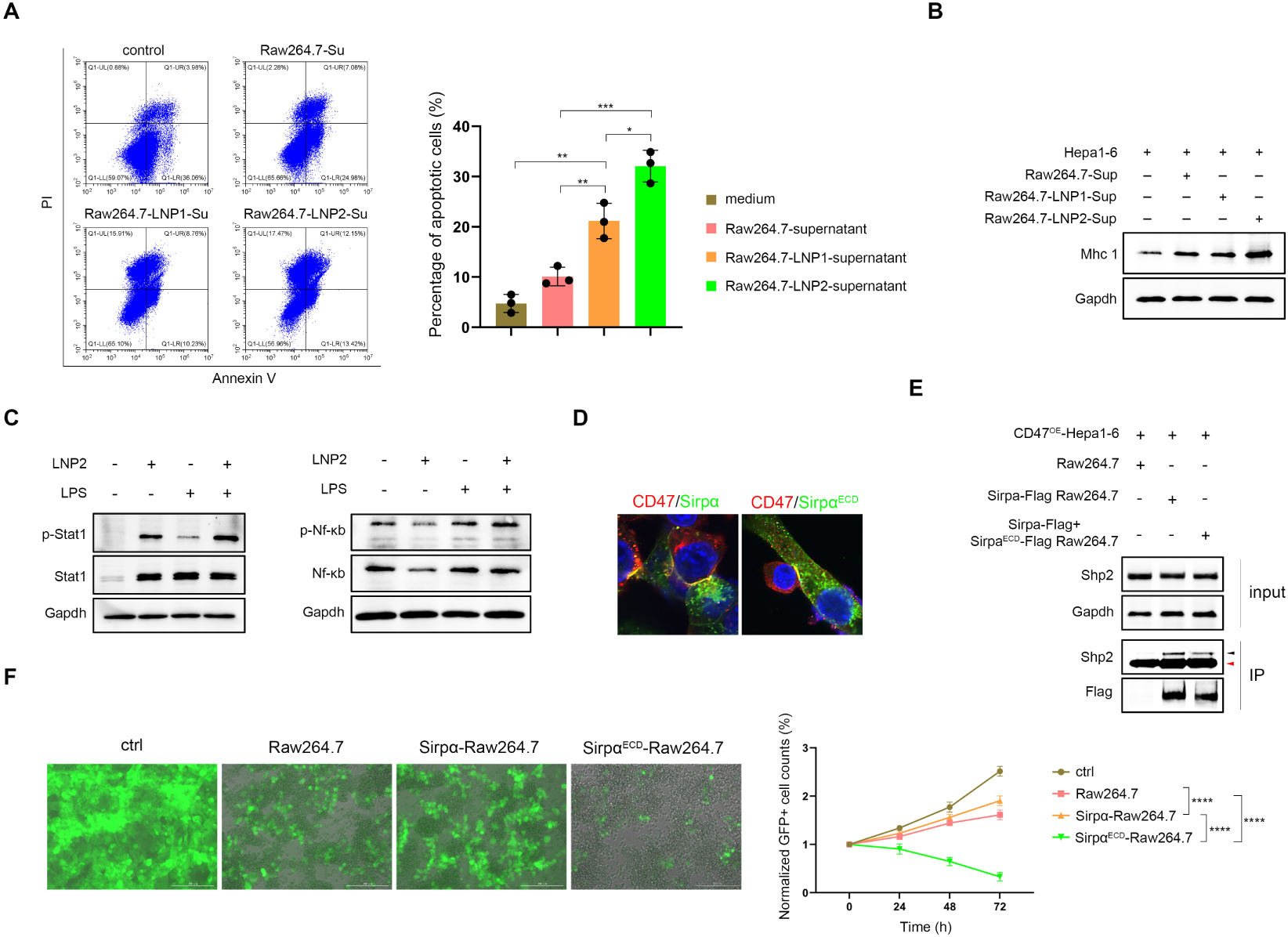
Ifn-γ and Sirpα^ECD^ proteins enhance CAR-M anti-tumor activity through distinct mechanisms. (A) Apoptosis analysis of Hepa1-6 cells exposed to different culture supernatants. Supernatants were collected from Raw264.7 or CAR-M cells 48 hours post-LNP1/2 transfection, filtered to remove live cells, and used to replace the culture medium of Hepa1-6 cells. Apoptosis was assessed 24 hours later using Annexin V and PI staining, with the right panel showing the proportion of PI^+^ cells across groups. (B) Western blot analysis of MHC I protein expression in Hepa1-6 cells. Tumor cells were pre-treated for 24 hours with supernatants from Raw264.7 or CAR-M cells before protein extraction. (C) Western blot detection of p-Stat1/Stat1 or p-Nf-κb/Nf-κb expression levels in Raw264.7 cells following 24-hour incubation with LNP2 or LPS. (D) Immunofluorescence analysis of Cd47-Sirpα or Cd47-Sirpα^ECD^ co-localization. To avoid interference from endogenous Cd47 and Sirpα, Cd47-Myc was overexpressed in Hepa1-6 cells, and Sirpα-Flag or Sirpα^ECD^-Flag was overexpressed in Raw264.7 cells. Following a 4-hour co-culture, Myc and Flag antibodies were used to assess the interaction between Cd47 and Sirpα/Sirpα^ECD^. (E) Immunoprecipitation assay and western blot examination of Shp2-Sirpα protein interaction in Raw264.7 cells with Sirpα or Sirpα^ECD^ overexpression. The black arrow indicates the anti-Shp2 band, whereas the red arrow marks the heavy chain. Cd47, Sirpα or Sirpα^ECD^ were all overexpressed in tumor or macrophage to highlight Cd47-Sirpα interaction axis. To mimic the negative dominance effect of Sirpα^ECD^ in CAR-M, both Sirpα-Flag and Sirpα^ECD^-Flag vector were co-lentiviral transfected into Raw264.7 cells. After 6 hours co-culture with CD47^OE^-Hepa1-6 cells, anti-Flag antibody was used for immunoprecipitation assay. The same amount of precipitated protein was loaded for western blot examination, in comparison with Sirpα-Flag overexpression group. (F) Live-cell imaging of GFP-expressing Hepa1-6 cells after 72 hours of co-culture with Raw264.7 cells overexpressing Sirpα or Sirpα^ECD^ at a 1:1 E:T ratio, along with time-course quantification of GFP^+^ cells. The left panel displays the co-culture status at 72 hours, while the right panel records the dynamic changes in GFP^+^ cell counts over time. The data were presented as mean ± s.e.m., and two-way ANOVA analysis was used to calculate the significance. \*\*\*\**p* < 0.0001.

The CD47–SIRPα signaling axis is well established for its role in protecting CD47-expressing tumor cells from phagocytosis by SIRPα-expressing macrophages. Upon ligand binding, the ITIM of cytoplasmic SIRPα recruits phosphatases SHP1/2, thereby inhibiting phagocytosis. In our design, Sirpα^ECD^ is engineered to competitively bind Cd47 to suppress endogenous Sirpα–Shp2 signaling in macrophages. To validate this, we first confirmed that the truncated Sirpα protein, Sirpα^ECD^, was localized at the cell membrane of CAR-M (Fig. 3D). Furthermore, immunofluorescence analysis demonstrated that Sirpα^ECD^ could bind the Cd47 ligand in a manner similar to the native Cd47–Sirpα interaction during macrophage and tumor cell co-culture (Fig. 3D). Third, we assessed Shp2 abundance in Flag antibody-precipitated proteins from macrophages overexpressing Sirpα^ECD^-Flag and Sirpα-Flag, and found that Shp2 enrichment was significantly reduced upon Sirpα^ECD^ co-expression (Fig. 3E), confirming that Sirpα^ECD^ disrupts the interaction between ITIM and Shp2 via a negative dominant manner. Finally, to determine the extent to which Sirpα^ECD^-mediated Shp2 inhibition contributes to tumor cell killing, we co-cultured Hepa1-6 tumor cells with macrophages with or without Sirpα^ECD^ overexpression. Our results showed that Sirpα^ECD^ significantly enhanced the ability of macrophages to eliminate tumor cells (Fig. 3F).

Overall, the Ifn-γ and Sirpα^ECD^ incorporated into our CAR structure serve distinct yet complementary functions. Ifn-γ activates macrophages and enhances tumor antigen presentation, while Sirpα^ECD^ strengthens macrophage phagocytic activity. Given the critical roles of macrophages in antigen presentation and modulation of the immune microenvironment, the co-expression of Ifn-γ and Sirpα^ECD^ in CAR-M may synergistically enhance its anti-tumor efficacy.

### LNP-mRNA in vivo delivery efficiently suppress orthotopic liver tumor growth in a macrophage-dependent manner

Next, we evaluated the in vivo anti-tumor activity of CAR-M. Previous studies have shown that the liver is the primary site for intravenously injected LNPs [25, 26]. Fluorescent signal distribution of LNP-encapsulated luciferase mRNA in mice further confirmed this (Fig. S6A). Moreover, LNPs encapsulating GFP mRNA revealed that Kupffer cells, a type of tissue-resident macrophage, internalized most of the LNP (Fig. S6B, C), consistent with previous reports [27]. We then established a liver orthotopic xenograft tumor model in mice using Hepa1-6 cells overexpressing human GPC3. LNPs or drug was administered intravenously at regular intervals, and tumor fluorescence was dynamically monitored (Fig. 4A). We firstly asked whether LNP2 would cure the orthotopic tumor, and whether this effect was dependent on in situ macrophages or not. After two doses administration, it revealed that LNP2 treatment significantly alleviated the tumor burden, while clodronate liposomes, an in vivo macrophage depletion drug, abolished the tumor regression effect (Fig. 4B, Fig. S7A), indicating the anti-tumor activity of LNP2 was dependent on endogenous macrophages.

**Figure 4.**
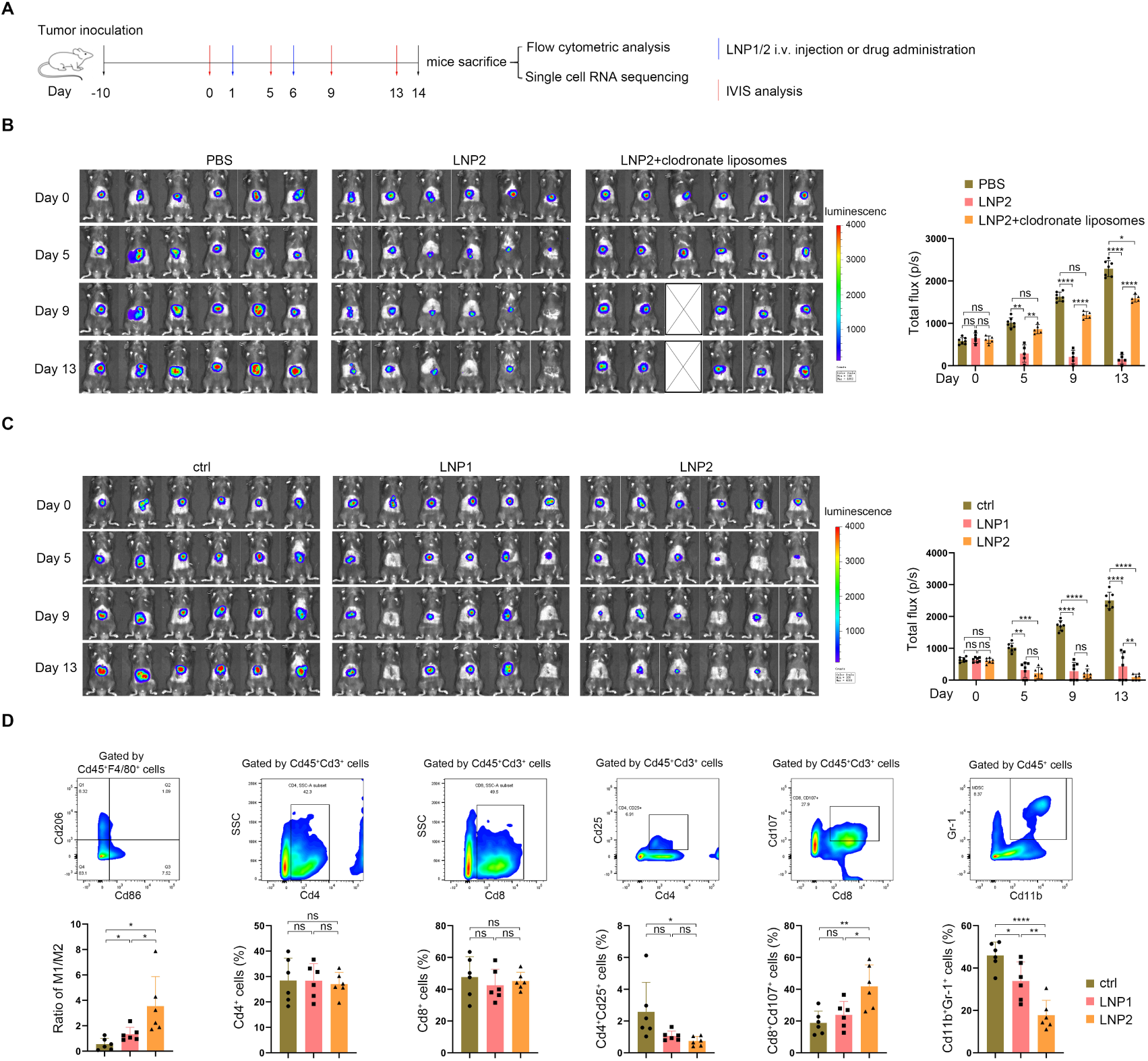
In situ generated CAR-Ms inhibit tumor progression in a mouse liver orthotopic xenograft model. (A) Schematic illustration of the experimental design to evaluate anti-tumor activity of in vivo-delivered LNP1/2 in a liver orthotopic xenograft tumor model. (B, C) In vivo bioluminescence imaging showing time-dependent tumor growth in mice following treatment with LNPs or clodronate liposomes. The right panel presents the quantification of bioluminescence signal across different treatment groups over time. (D) Flow cytometric analysis of the immune cell composition in the liver tissue surrounding the xenograft tumor. The upper panel illustrates the gating strategy for different immune cell populations, while the lower panel displays the quantified proportions of immune cells within the TME. Data were presented as mean ± s.e.m., and significance was evaluated by t-test analysis. \**p* < 0.05; \*\**p* < 0.01; \*\*\**p* < 0.001; \*\*\*\**p* < 0.0001. ns, not significant.

We then compared the anti-tumor efficiency of LNP1 and LNP2 in liver orthotopic xenograft tumor model. After two doses administration, both LNP1- and LNP2-treated mice exhibited significant tumor regression, with LNP2 showing superior efficacy compared to LNP1 (Fig. 4C, Fig. S7B). Then the mice were sacrificed and liver tissues were harvested for further analysis. We first examined the immune microenvironment within the tumor region. Flow cytometric analysis revealed a significant increase in the ratio of Cd86^+^ to Cd206^+^ macrophages following LNP2 treatment (Fig. 4D). Moreover, the proportions of Cd4^+^ or Cd8^+^ T cells remained unchanged after LNP treatment; however, regulatory T cells (Tregs) and cytotoxic Cd8^+^ T cells exhibited apparent responses to LNP treatment (Fig. 4D), suggesting that the in situ generated CAR-M contributed to the establishment of a pro-inflammatory environment at the expense of immunosuppressive components within the TME. Additionally, the population of myeloid-derived suppressor cells (MDSCs) was restricted to a low level (Fig. 4D). Next, immunohistochemical (IHC) analysis of liver tissues showed that the expression of *p*-Stat1, a downstream effector of Ifn-γ, was markedly elevated in liver tissue cells in LNP2 treatment group (Fig. S7C, D).

### In situ generated CAR-M remodels the TME into a pro-inflammatory and anti-tumor state

To further evaluate the immune microenvironment following LNP treatment, cells surrounding the solid tumor (or scar tissues in LNP1 or LNP2 treatment groups) within TME were carefully collected for single-cell RNA sequencing (scRNA-seq) analysis (Fig. 4A). After data filtering and quality control, a total of 11628, 13274 or 14968 cells were analyzed from the control, LNP1, or LNP2 treatment groups, respectively. Cell annotation and t-distributed stochastic neighbor embedding (t-SNE) analysis identified nine major cell populations across the treatment groups, with varying proportions observed among the groups (Fig. 5A, B). The scRNA-seq results indicated that monocytes, neutrophils, T/NK cells, fibroblasts, and endothelial cells play key roles in modulating the inflammatory status within the TME (Fig. 5C). Macrophages primarily reshape the immune microenvironment by secreting inflammatory cytokines and presenting antigens to T cells [5]. As a feedback response to Ifn-γ produced by CAR-M, the expression of *B2m*, which encodes a subunit of Mhc-I, was significantly elevated in the LNP2 treatment group (Fig. S8A). To determine whether LNP2-mediated in situ programming of CAR-M activated anti-tumor immunity within the TME, we further analyzed specific cell subsets.

**Figure 5.**
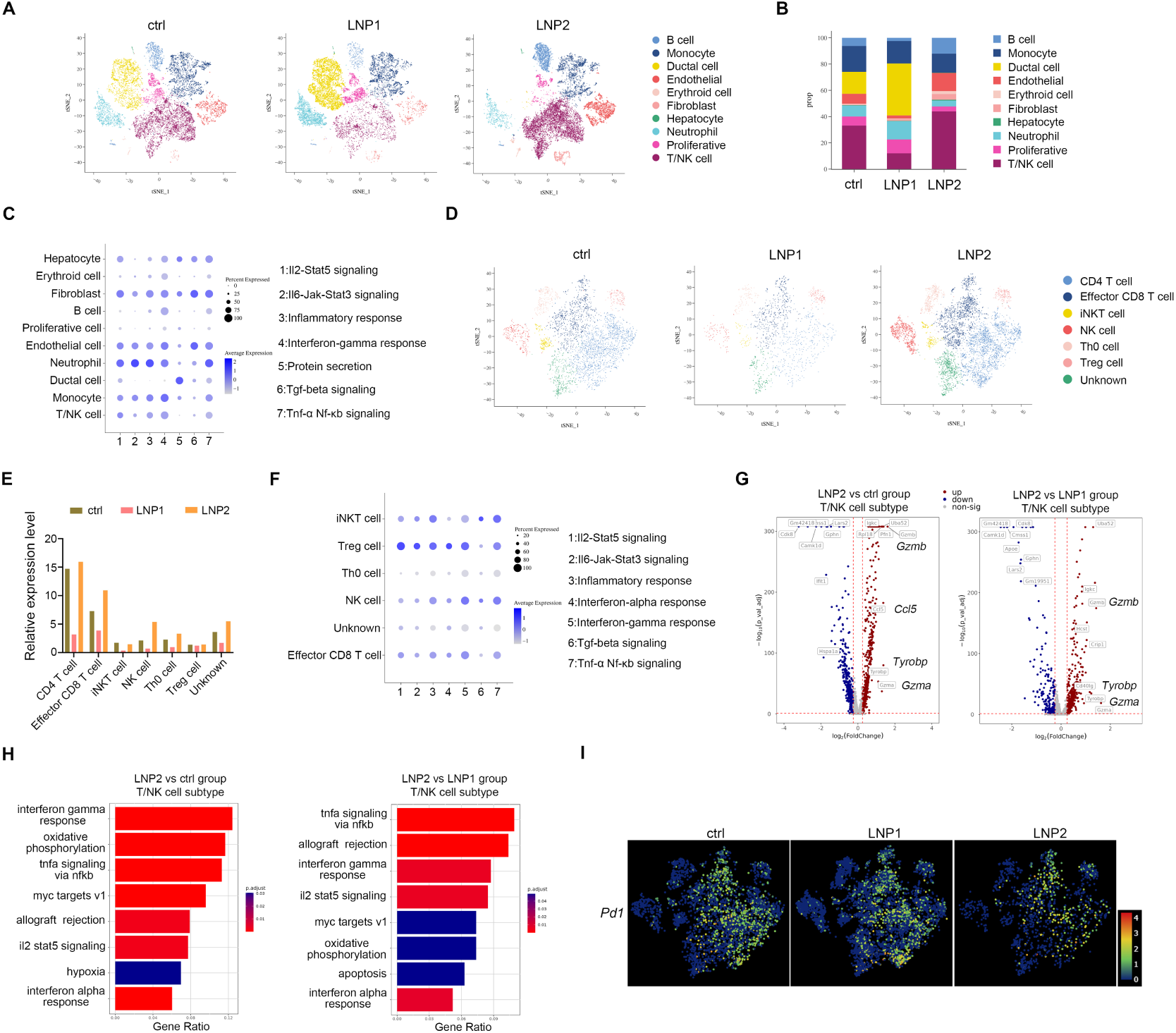
Single-cell RNA sequencing reveals that LNP2-armed CAR-Ms promote an anti-tumor immune environment in the mouse liver. (A) t-SNE visualization of scRNA-seq data from different samples. Cells were clustered into nine distinct groups, with color codes indicating cell identities. (B) Stacked bar chart showing the relative proportions of the nine cell clusters across samples. (C) Bubble plot illustrating the activity of pro-inflammatory signaling pathways in different cell clusters. The size and color intensity of each bubble reflect the relevance and significance of the pathway activity. (D) t-SNE plot depicting the composition of T/NK cell subsets across different samples, with color-coded cell identities. (E) Quantitative analysis of the relative proportions of various T/NK cell subpopulations in each sample. (F) Bubble plot showing the activity of pro-inflammatory signaling pathways within different T/NK cell subsets. (G) Volcano plots highlighting the most significantly enriched genes in T/NK cells from LNP2-treated mice compared to ctrl or LNP1-treated mice. (H) Bar plot displaying the most significantly enriched Hallmark gene sets differentially expressed between LNP2 and ctrl or between LNP2 and LNP1 groups. Hallmark gene sets were obtained from the MSigDB database. (I) Pd1 mRNA expression in T/NK cells across different samples, with color intensity reflecting gene expression levels.

t-SNE analysis revealed that dendritic cells, Kupffer cells, macrophages, and certain granulocytes constituted the monocyte cell subset in the TME (Fig. S8B). We first compared the top-expressed genes in monocytes across the treatment groups. Heatmap results showed that pro-inflammatory genes such as *Lyz* and *Nos2* were significantly enriched in macrophages following LNP2 treatment (Fig. S8C). Moreover, cytokine-mediated pro-inflammatory signaling pathways, particularly the Ifn-γ response, were highly activated in the monocyte subset in the LNP2 treatment group (Fig. S8D), confirming that LNP2-armed CAR-M contributes to the establishment of a pro-inflammatory microenvironment.

Next, we evaluated the T/NK cell subset, which mainly included Cd4+ T cells, effector Cd8+ T cells (Teff), iNKT cells, NK cells, naive T cells (Th0), and Treg cells (Fig. 5D). Further analysis revealed that the proportions of Teff, iNKT, and NK cells were increased in the LNP2 treatment group, and these cells were associated with pro-inflammatory signaling processes in the TME (Fig. 5E, F). Comparative analysis of the top-expressed genes in the T/NK subset across treatment groups showed that cytotoxic factors such as *Gzma/b*, chemokines like *Ccl5*, and the immune signaling adaptor gene *Tyrobp* were significantly enriched in the LNP2 group (Fig. 5G). Pathway analysis further confirmed that LNP2-induced CAR-M promotes a pro-inflammatory and cytotoxic immune status (Fig. 5H). Subsequently, we examined the expression levels of immune exhaustion markers, including *Pd1*, *Lag3*, and *Tim3*, and found that all three were markedly downregulated in the LNP2 treatment group (Fig. 5I, Fig. S8E). In contrast, effector memory T cell markers were upregulated in the LNP2 group (Fig. S8F), suggesting effective antigen cross-presentation between CAR-M and T cells within the TME.

Finally, we assessed the anti-tumor efficacy of memory T cells against GPC3-expressing Hepa1-6 tumor cells using a tumor re-challenge assay (Fig. S9A). Twenty-six days post-tumor inoculation, control mice reached the euthanasia criteria, whereas mice previously cured by LNP2 administration effectively rejected tumor re-challenge (Fig. S9B), demonstrating that in situ generation of CAR-M via LNP2 delivery facilitates the formation of durable immune memory within the TME.

Collectively, these data indicate that both LNP1 and LNP2 treatments remodel the tumor immune microenvironment, with LNP2 being more effective in establishing a pro-inflammatory niche that supports long-term anti-tumor immunity.

### Establishment of the co-culture system using human PBMC derived CAR-M and liver caner organoid

To evaluate the translational potential of our CAR design for human applications, we synthesized a novel CAR mRNA encoding human IFN-γ and the extracellular domain of human SIRPα, which was encapsulated into LNPs. Human mononuclear leukemic cell line THP1 and PBMC-derived macrophages were induced into the M1 phenotype following treatment with PMA, IFN-γ, and LPS [28]. Notably, CAR-M generated via LNP delivery exhibited a more activated phenotype, as evidenced by the upregulation of pro-inflammatory genes and increased polarization toward the M1 phenotype (Fig. S10A, B). Furthermore, using a human HCC cell line Huh7, which highly expresses the GPC3 antigen, we demonstrated that CAR-M exerts potent anti-tumor activity in co-culture assays (Fig. S10C), consistent with findings from murine models.

To further validate the ability of CAR-M to recognize and eliminate endogenous tumor cells, we employed a human liver organoid system. Liver tissues were obtained from clinical samples and cultured according to previously described protocols [29]. Compared with healthy liver organoids (HLO), liver cancer organoids (LCO) exhibited significantly higher GPC3 expression levels (Fig. 6A, B). To evaluate the direct anti-tumor effects of CAR-M, we established a macrophage-organoid co-culture system in which HLO or LCO were incubated with M1-polarized macrophages or CAR-M derived from PBMC or THP1, respectively (Fig. 6C, Fig. S10D). Organoid morphology was monitored every 24 hours, and the results showed that only LCO co-cultured with CAR-M exhibited evident disaggregation or disruption (Fig. 6C, D). Moreover, cell viability assays confirmed that CAR-M specifically targeted and destroyed tumor cells (Fig. 6E). Collectively, these findings demonstrate that our CAR-M strategy enables the specific recognition and elimination of GPC3^+^ tumor cells with minimal off-target effects on healthy cells, highlighting its promising clinical application potential.

**Figure 6.**
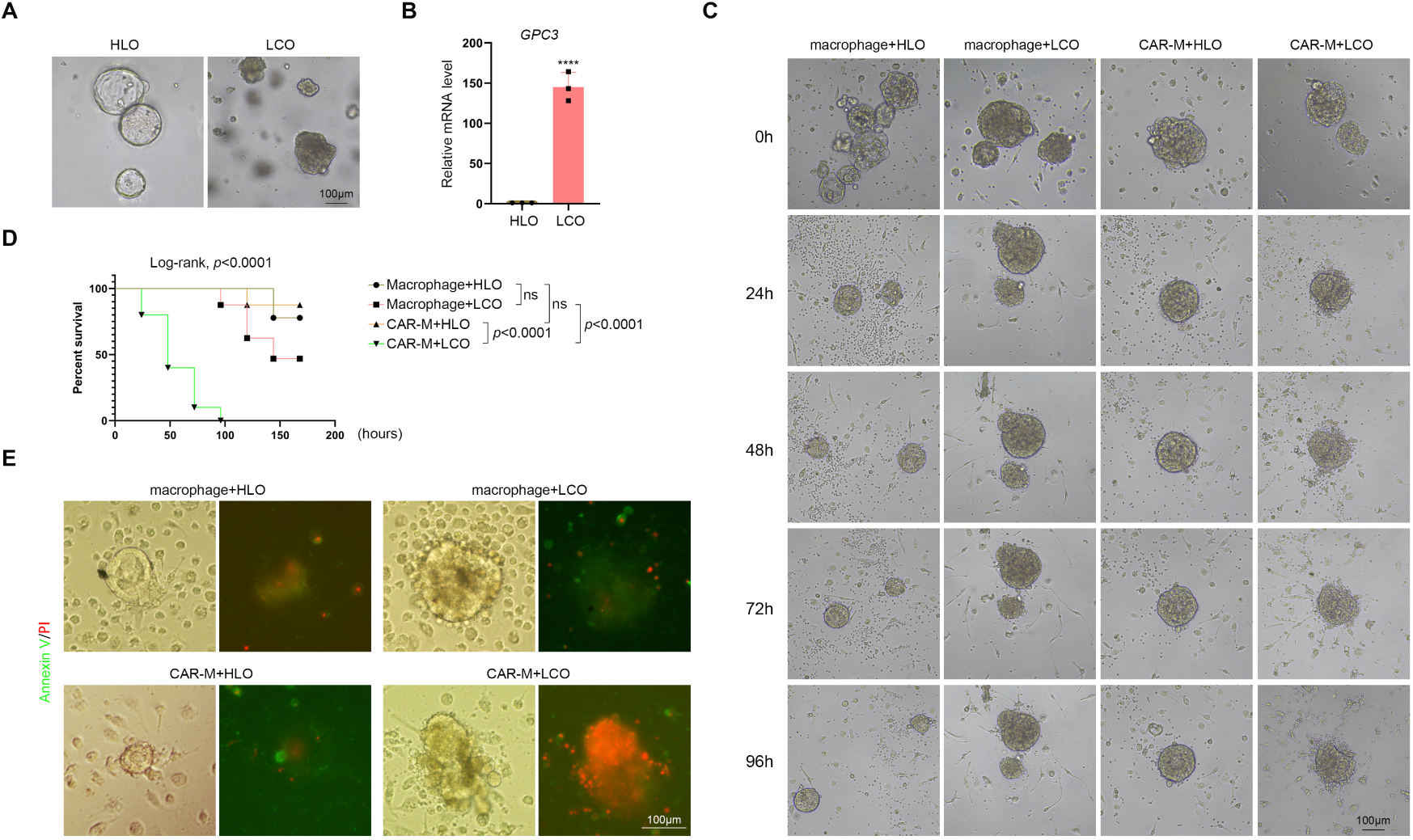
PBMC derived CAR-M specifically eliminates liver cancer organoid in vitro. (A) Establishment and morphology of HLO or LCO derived from clinical samples. (B) qRT-PCR detection of GPC3 mRNA expression in HLO and LCO. The organoids used for qRT-PCR were established from three different patient samples. (C) Morphological dynamics of HLO and LCO co-cultured with PBMC or PBMC-derived CAR-M cells over time. The organoids shown in the image are representative of at least 30 organoids per group. (D) Kaplan-Meier survival analysis of HLO and LCO after long-term co-culture with macrophages or CAR-Ms. The co-culture system was observed and recorded every 24 hours. Organoids exhibiting apparent disaggregation or disruption were considered as death events. (E) Fluorescence images showing the Annexin V and PI staining in HLO or LCO after 72 hours co-culture with macrophages or CAR-Ms. The organoids shown in the image are representative of at least 20 organoids per group.

## Discussion

Owing to their tumor infiltration capability, inherent plasticity, and dual roles in inflammation and immune modulation, macrophages are considered promising CAR-engineered immune cell carriers for combating solid tumors. Ideally, infiltrated CAR-Ms are expected to fulfill the following functions: direct phagocytosis of tumor cells, sustained M1 polarization to secrete pro-inflammatory factors that remodel the TME, and antigen presentation to other immune cells. Therefore, optimizing the CAR construct to achieve these functions remains a central focus in CAR-M research. The LPS-TLR4 and IFN-γ-IFNGR signaling axes are two classical pathways that induce macrophage M1 polarization. Previous studies have shown that CAR constructs incorporating constitutively active TLR4 signaling outperform first-generation CAR-Ms containing only CD3ζ [14]. CAR-Ms incorporating IFNGR or functional elements of IFN-γ have also been reported to exhibit potent anti-tumor activity, with the latter showing particularly strong efficacy [30, 31]. In our study, we treated CAR-Ms with LPS to maintain TLR4/NF-κB activation and found that TLR4 activation enhanced the anti-tumor efficacy of LNP1-armed CAR-Ms, consistent with prior reports. Meanwhile, LNP2-armed CAR-Ms exhibited active p-STAT signaling and demonstrated comparable or even superior anti-tumor activity to the LPS-treated group. Furthermore, our in vitro and in vivo data revealed a durable pro-inflammatory state and anti-tumor immune microenvironment established by CAR-Ms, likely due to the self-sustaining effect of IFN-γ. Considering the marked upregulation of MHC-I protein expression and direct cytotoxic effects on tumor cells, IFN-γ may represent a more advantageous strategy than TLR4 activation for promoting CAR-M persistence in CAR construct design.

A critical challenge in CAR-M therapy is how to effectively abrogate the CD47-SIRPα signaling axis. Previous strategies, such as incorporating a secretory CD47-blocking antibody into the CAR construct [32], or using shRNA to silence SIRPα in CAR-Ms [33], have demonstrated notable anti-tumor activity in solid tumors. However, CD47 is a broadly expressed ligand across various cell types, and its systemic blockade may lead to non-specific macrophage-mediated cytotoxicity, potentially causing severe adverse effects during CAR-M therapy. Previous clinic trial using CD47 monoclonal antibody in solid tumors have reported frequent toxicities, including transient anemia, fatigue, headache, and lymphopenia in most treated patients [34]. Furthermore, proteins generally exhibit longer half-lives than RNA molecules [35, 36], suggesting that our dominant-negative SIRPα^ECD^ protein-mediated inhibition of SIRPα signaling may provide a more sustained effect compared to transient shRNA knockdown. Indeed, our in vivo results showed detectable levels of SIRPα^ECD^ protein in in situ-programmed macrophages even after tumor regression. Additionally, beyond its interaction with SIRPα, CD47 also binds to integrins and thrombospondin-1, contributing to diverse cellular processes such as migration, proliferation, and adhesion [37]. By consuming available CD47 through direct occupancy via SIRPα^ECD^, our dominant-negative blockade may offer a more comprehensive and effective inhibition strategy than SIRPα silencing alone. Collectively, our in vivo delivery and macrophage reprogramming approach presents a promising and potentially superior solution for overcoming the CD47–SIRPα immune checkpoint in CAR-M therapy.

To date, all FDA-approved CAR-immune cell therapies rely on ex vivo engineering of autologous T cells. This personalized manufacturing process involves isolating the patient’s T cells, genetically modifying them using lentiviral or retroviral vectors, administering lymphodepleting regimens, and reinfusing the engineered CAR-T cells into the same patient [38]. The complexity and high cost of this process limit its broad clinical application. Although allogeneic T cells from healthy donors offer an off-the-shelf alternative, they require additional genetic modifications to eliminate αβ T cell receptors (TCRs) and carry a risk of life-threatening graft-versus-host disease (GvHD) [39]. In contrast, in vivo CAR immune cell engineering strategies offer a promising alternative to overcome the limitations of both autologous and allogeneic approaches. Various in vivo delivery platforms have been developed to program T cells or macrophages, including LNPs, lentiviruses, adeno-associated viruses (AAVs), virus-mimetic fusogenic nanovesicles (FuNVs), and hybrid strategies involving bioinstructive implantable scaffolds [12]. Among these, LNPs are the most widely used, largely due to their successful application in mRNA vaccines against COVID-19. The transient nature of this in vivo approach minimizes potential toxicities and allows for repeated dosing. Recent studies have shown that all patients with relapsed or refractory multiple myeloma who received BCMA-targeting lentiviral vectors for in vivo T-cell engineering achieved partial or complete responses [40], highlighting the therapeutic potential of in vivo CAR-immune cell engineering. Given the liver-targeting properties of LNPs, LNP-mediated mRNA delivery for CAR programming represents a relatively low-cost, easily producible, and ready-to-use strategy, making it a promising approach for treating liver cancer. According to previous reports, HCC is characterized by a high infiltration of immunosuppressive myeloid cells, which create a pro-tumoral immune microenvironment and hinder the function of newly infiltrated CAR-T or CAR-M cells. From this perspective, in situ engineered CAR-M, which is generated through reprogramming TAM within the TME, is undoubtedly a more favorable strategy compared to adoptive CAR-M transfer.

To sum up, we developed a novel CAR design combined with an in vivo programming strategy to generate CAR-M for immunotherapy against HCC. Both in vitro and in vivo data indicated LNP encapsulated CAR mRNA could efficiently reprogram macrophages into a M1 pro-inflammatory phenotype. Benefited from the integrated IFN-γ and SIRPα^ECD^ functional modules, our CAR structure design enabled macrophages to eliminate tumor cells and remodel the tumor microenvironment like a vaccine in a more effective and persistent manner compared to previous CAR-M approaches, suggesting a synergistic working model. Importantly, our CAR-M specifically recognized and eradicated human liver tumor organoid while sparing healthy organoids, thereby validating its strong translational potential for clinical applications.

## Methods

### Cell Lines

Mouse Raw264.7, Hepa1-6 cell lines, and human THP1, Huh7, HepG2 and 293T cells lines are all purchased from the Culture Collection of Chinese Academy of Sciences (Shanghai, China), and are operated according to the guideline. Generally, cells were cultured in DMEM or RPMI 1640 (Thermo) medium containing 10% fetal bovine serum (FBS, Vazyme), 1% penicillin/streptomycin (Thermo) and a 5% CO_2_ atmosphere at 37 ℃. BMDM was harvested from mice femurs and cultured in DMEM medium with 10% FBS, 1% penicillin/streptomycin and 10 ng/ml mouse Macrophage colony-stimulating factor (M-CSF, Sino Biological, 51112-M08H). PBMC was purchased from Milecell Bio Inc., and was cultured in DMEM medium with 10% FBS, 1% penicillin/streptomycin and 50 ng/ml human M-CSF (Yeasen, 91103ES). Before LNP-mRNA transfection, Raw264.7 or THP1 cells were stimulated into M1 stage using 100 ng/ml Phorbol 12-myristate 13-acetate (PMA) (Sigma-Aldrich, P8139), 100 ng/ml LPS (Sigma-Aldrich, L4391) and 20 ng/ml IFN-γ (Sino Biological, 11725-HNAS). 500 ng/ml LNP-mRNA was incubated with 1×10^6^ cells to generate CAR-M for the following experiments.

### LNP-mRNA preparation

The CAR mRNAs are composed of anti-human GPC3 scFv, a human or mouse CD8α hinger, a human or mouse CD28 transmembrane and co-stimulatory domain, a human or mouse CD3ζ immunoreceptor tyrosine-based activation motif (ITAM), and human or mouse tandem IFN-γ-2A-SIRPα^ECD^ coding sequence. The mRNAs were synthesized and encapsulated into LNP by Magicrna Inc. according to previous reports [41]. Briefly, mRNAs were in vitro transcribed and the UTPs were replaced with one-methyl pseudouridine (m1Ψ)-5′-triphosphate (TriLink) to enhance the stability. After m7G capping and purification, the mRNAs quality was assessed following the manufacturer kit (DNF-472-1000). Next, the newly synthesized lipids and mRNAs were mixed in a microfluidic chip device with a 5:1 N/P ratio. After dialysis, the end products were diluted in 1×PBS containing 20×10^−3^ M Tris with 7.4 pH value, and were aliquoted and stored at −80°C prior to use. For LNP-mRNA in vivo distribution assay, 5μg luciferase or eGFP mRNA encapsulated by LNP (Magicrna) was intravenously injected into mice. About 6 hours later, mice was sacrificed and fluorescent signals was monitored in the main organs accordingly.

### Flow cytometry

The cultured cell lines or mice liver tissues were digested into single cell suspension, and incubated with the fluorescence tagged antibodies listed as follows, Cd3 (BD Pharmingen, 561827), Cd4 (BD Pharmingen, 552775), Cd8 (BD Pharmingen, 566409), Cd11b (BD Pharmingen, 561688), Cd86 (Biolegend, 105115), Cd25 (BD Pharmingen, 562606), Cd45 (BD Pharmingen, 561037), Cd107a (BD Pharmingen, 560646), Cd206 (Biolegend, 141707), F4/80 (BD Pharmingen, 565411), Gr-1 (BD Pharmingen, 553128), Sirpα (Invitrogen, 17-1729-42), CD86 (Biolegend, 374207), and CD206 (Biolegend, 321105). Then, the cells were examined by CytoFLEX Flow Cytometer (Beckman) and analyzed by CytExpert or Flowjo software thereafter.

### Immunofluorescence

The freshly dissected mice liver tissues were fixed by paraformaldehyde for 24 hours. After paraffin embedding, the samples were sectioned at intervals of 3μm by paraffin microtome (Leica Biosystems). After deparaffinization and rehydration, the sectioned samples were subjected to Hematoxylin and Eosin (H&E) staining, or were incubated with the following antibodies respectively, Mhc-I (Beyotime, AG2146, 1:1000), PD1 (Proteintech, 66220-1-Ig, 1:200). The samples were then reacted with the corresponding secondary fluorescent antibodies (1:500, A-21206, A-21203, Invitrogen) and DAPI staining (Beyotime). Fluorescent images were captured by confocal microscope System (Nikon, AX).

### Western blot and immunoprecipitation

The protein were extracted from cells with different treatment using radio-immunoprecipitation assay lysis buffer (RIPA, Beyotime). After evaluating protein concentration by BCA assay (Thermo), proteins of the same mass were loaded and separated using sodium dodecyl sulfate-polyacrylamide gel electrophoresis and transferred onto polyvinylidene difluoride membranes (Millipore). The membranes were then incubated with different antibodies at 4°C overnight. The antibodies were listed as follows: anti-Flag (Proteintech, 66008-4-Ig, 1:500), anti-Myc antibodies (Proteintech, 16286-1-AP, 1:500), anti-GPC3 (Proteintech, 30021-1-AP, 1:1000), anti-GAPDH (Proteintech, 60004-1-Ig, 1:50000), anti-Nf-κb p65 (Selleck, F0006, 1:1000), anti-phospho-Nf-κb p65 (Ser536) (Selleck, F0155, 1:1000), anti-Stat1 (Selleck, F0263, 1:1000), anti-phospho-Stat1 (Ser727) (Selleck, F0451, 1:1000). Next, the membranes were incubated with species-matching secondary antibodies, and was captured using chemiluminescence by gel imaging system (ChemiDoc MP Imaging System, Bio-Rad). For immunoprecipitation, the extracted proteins were firstly incubated with anti-Flag (Proteintech, 66008-4-Ig, 1:20000) or anti-Myc antibodies (Proteintech, 16286-1-AP, 1:5000) at 4°C overnight on a rotary shaker, and then protein A/G Magnetic Beads (Vazyme, PB101) were supplemented and incubated at room temperature for 2 hours. After enrichment by magnetic frame, the protein samples were loaded into gel for Western blot detection.

### Mass spectrometry

LNP1 or LNP2 armed BMDM cells were co-cultured with GPC3 overexpressing Hepa1-6 cells respectively in serum-free medium. After 24 hours co-culture, the supernatant was harvested for Mass spectrometry (MetwareBio). In brief, the protein was purified and quantified by BCA assay kit. After proteolytic desalting, the peptide concentration was determined using a Pierce™ Quantitative Peptide Assay Kit with standards (Thermo Fisher). Next, the samples were separated using the Vanquish Neo UHPLC liquid chromatography system, and were subjected to Data-Independent Acquisition mass spectrometry analysis using the Orbitrap Astral high-resolution mass spectrometer (Thermo Scientific). Finally, the raw data were analyzed using DIA-NN(v1.8.1) with library-free method.

### Live-cell imaging

About 5×10^4^ GFP marked Hepa1-6 tumor cells with or without GPC3 overexpression were cultured in 48-well plate with three replacates in each group. Macrophages or CAR-M cells with the same number were added into the wells respectively for long-term co-culture. GFP dynamics in each well was monitored by live cell imaging system (Agilent BioTek, Lionheart FX) with the same working parameters at a regular time interval.

### Mice liver orthotopic xenograft tumor model

All animal experiments are carried out after obtaining authorization from the Animal Ethics Committee from Guangzhou Medical University (Approval No. GY2025-125). The mouse liver orthotopic xenograft tumor model was established according to a previously reported protocol [42]. Briefly, 4-week-old Balb/c mice were anesthetized via intraperitoneal injection of 1% pentobarbital sodium. Following laparotomy, approximately 1.3 million GPC3- and luciferase-overexpressing Hepa1-6 cells in a 20 μL volume were directly injected into the liver parenchyma. Subsequently, the abdominal cavity was sutured and the mice received an injection of penicillin-streptomycin to prevent infection. The following day, mice were monitored for any signs of distress or abnormalities. For bioluminescence imaging, mice were intraperitoneally injected with 150 mg/kg luciferin, anesthetized with isoflurane, and tumor signals were detected using the IVIS Lumina Series III (PerkinElmer) imaging system. For LNP administration, 5 μg of LNP-mRNA diluted in 200 μL sterile PBS was intravenously injected into each mouse.

### Single cell RNA sequencing

Mice with orthotopic tumors were sacrificed and the liver tissues were dissected. Then the tumor nodules were separated carefully and the TME cells were chopped and stored in tissue preservation solution provided by Seekgene Inc.. After removing erythrocytes, debris and dead cells, the cell nucleus was isolated using a Nucleus Isolation Kit (SHBIO, 52009-10). Single-cell RNA-Seq libraries were prepared using SeekOne® Digital Droplet Single Cell 3’ library preparation kit (SeekGene, Catalog No. K00202). The libraries were then sequenced on GeneMind SURFSeq 5000 with PE150 read length. The data was analyzed by seeksoul online server.

### Liver organoid

The study was approved by the Human Research Ethics Committee of the Second Affiliated Hospital of Kunming Medical University (Approval No. Shen-PJ-KE-2024-147). Written informed consent was obtained from all participants prior to the collection, research use, and publication of de-identified clinical data. Human healthy and tumor-derived liver organoids were established from clinical biopsy samples as previously described [29]. with the signed informed consent from patients. The organoids were embedded in growth factor-reduced Matrigel (Corning), and the culture medium was prepared according to the manufacturer’s instructions. Normal human liver organoids were cultured in advanced DMEM/F-12 medium (GIBCO, supplemented with HEPES, GlutaMax, and Penicillin-Streptomycin), and further supplemented with Bi-27 (MCE), N2 (MCE), 10 mM nicotinamide (MCE), 1.25 mM N-acetylcysteine (MCE), 10 nM gastrin (MCE), 50 ng/mL EGF (MCE), 100 ng/mL FGF10 (MCE), 100 ng/mL R-spondin1 (MCE), 50 ng/mL HGF (MCE), 50 ng/mL Wnt3a (MCE), 10 mM forskolin (MCE), 5 μM A83-01 (MCE), and 10 μM Rho Inhibitor γ-27632 (MCE). Tumor organoids were cultured in Hepatocellular Carcinoma Organoid Medium (bioGenous). During culture, the medium was refreshed every three days, and organoids were typically passaged every 7–10 days. For co-culture with macrophages, THP-1 cells or PBMC derived macrophages were transfected with LNP-encapsulated CAR mRNA. Organoids were dissociated into small cell clusters and embedded in Matrigel. Subsequently, Matrigel was removed using Organoid Isolation Solution (bioGenous) to collect the organoids. The isolated organoids were then added to 48-well plates with 7-10 organoids and 1×10^5^ macrophages or CAR-M in each well for co-culture.

### Statistical analysis

We used Graphpad Prism 8 software to analyze the statistical data and export the column charts. All in vitro data shown was representative of at least three independent experiments, whereas the mice in vivo data was repeated two times. The values were reported as the mean ± SD. Differences between the two groups were analyzed using the two-tailed unpaired Student’s t-test or two-way ANOVA analysis.

## Data availability Statement

The data in this study are available from the corresponding author upon reasonable request.

## Acknowledgments

This work was supported by The Seventh Affiliated Hospital, Sun Yat-sen University, with partial funding from the National Natural Science Foundation of China (Grants 82473133 and 82273089), the Xingdian Scholar Fund of Yunnan to Y.L.W., the Guangdong Basic and Applied Basic Research Foundation (Grants 2022A1515012232 and 2023A1515030027), and grants from projects 202501AS070120, 2024XKTDYS04, KLTIPT-2023-03, and YDYXJJ2024-0006.

## Author contributions

P.P.Z. conceived the whole project and designed the experiments with S.W., Y.L.W., and L.Z.; X.L., J.H., T.Z., and H.C.L., carried out all experiments and prepared figures; T.W. and Y.X.L. participated in the mice experiments; P.P.Z. drafted the manuscript; All authors read and approved the article.

---

**Figure S1.**
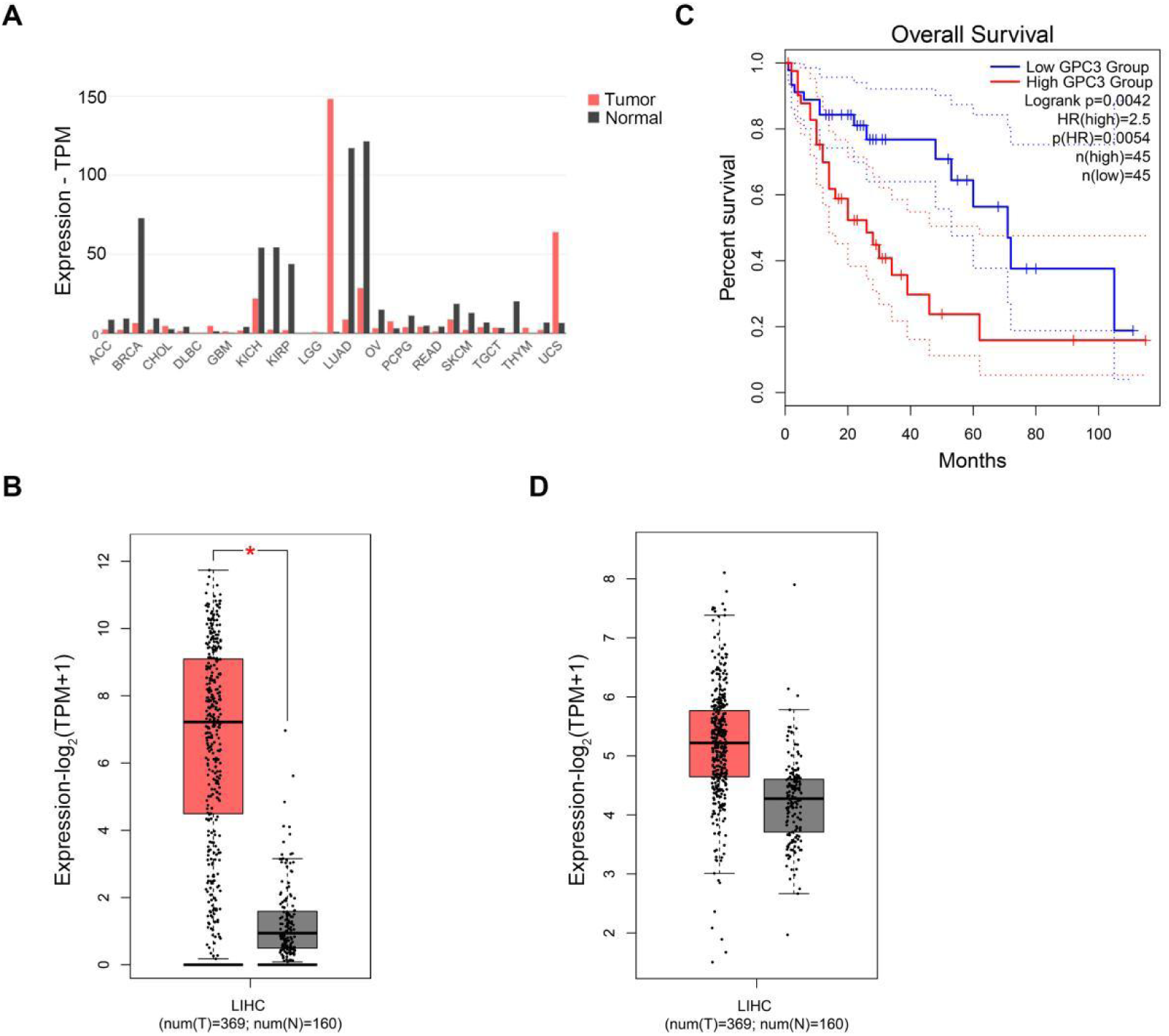
*GPC3* and *CD47* are highly expressed in liver hepatocellular carcinoma (LIHC). (A, B) *GPC3* is a relative specific biomarker highly expressing in LIHC. (C) High *GPC3* expression correlates with reduced overall survival in LIHC patients. (D) *CD47* is also highly expressed in LIHC patients. All data and images were obtained from the GEPIA online database.

**Figure S2.**
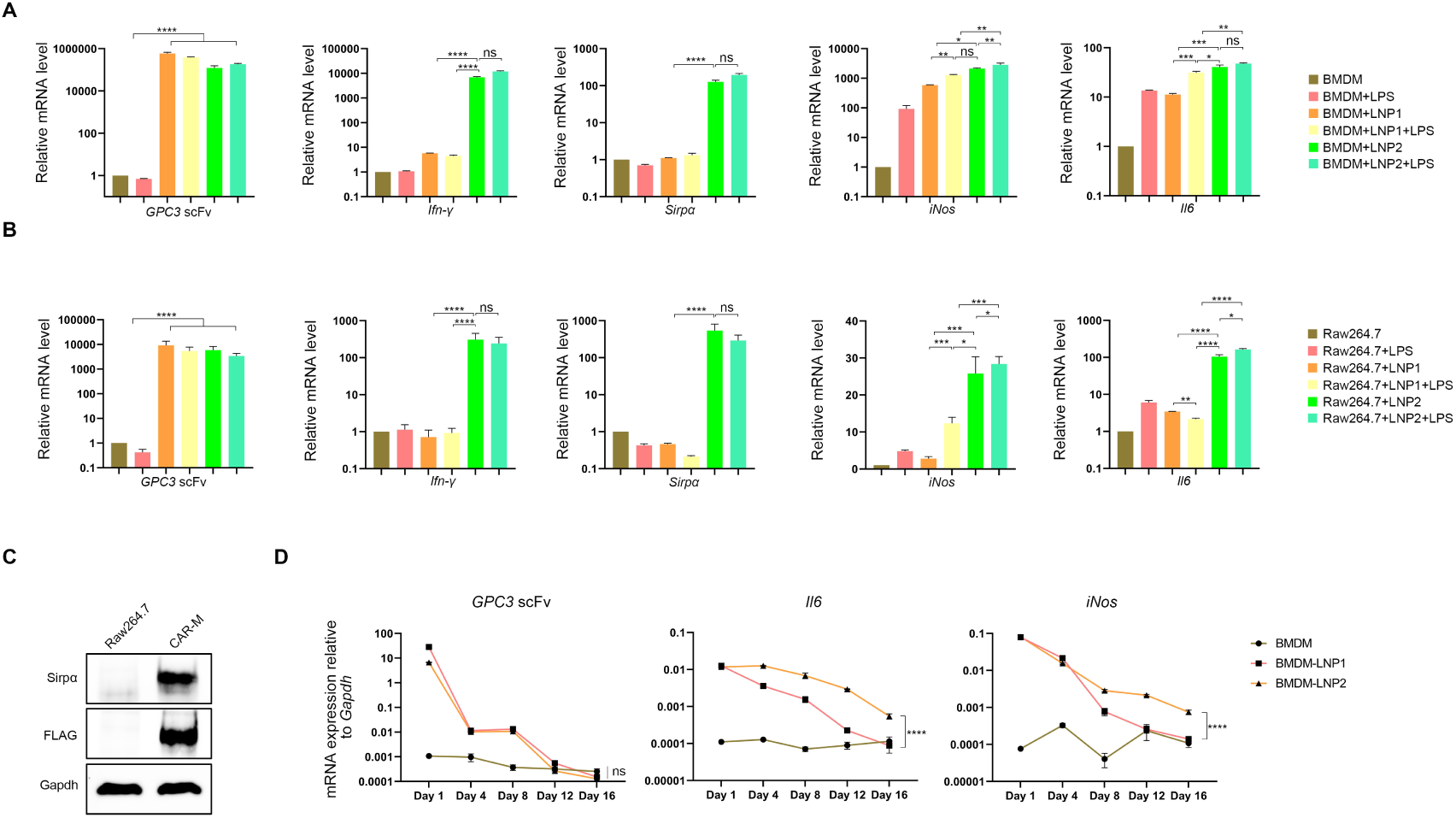
CAR-M cells armed with LNP1 or LNP2 exhibit an M1-polarized phenotype. (A, B) qRT-PCR analysis of CAR structural components and M1 macrophage marker gene expression in BMDM or Raw264.7 cells following treatment with LNP or LPS. Data are presented as mean ± s.e.m., and the significance was determined using two-tailed multiple t-tests analysis. \**p* < 0.05; \*\**p* < 0.01; \*\*\**p* < 0.001; \*\*\*\**p* < 0.0001. ns, not significant. (C) Western blot detection of Sirpα protein and Flag tag in Raw264.7 and CAR-M cells. (D) Statistical analysis of qRT-PCR results over time. Day 0 was defined as the day of LNP1/2 transfection in BMDMs, with cells collected at regular intervals up to day 16. The experiment was repeated three times. The Y-axis represents mRNA expression levels normalized to Gapdh. Statistical significance was calculated using two-way ANOVA analysis. \*\*\*\**p* < 0.0001. ns, not significant.

**Figure S3.**
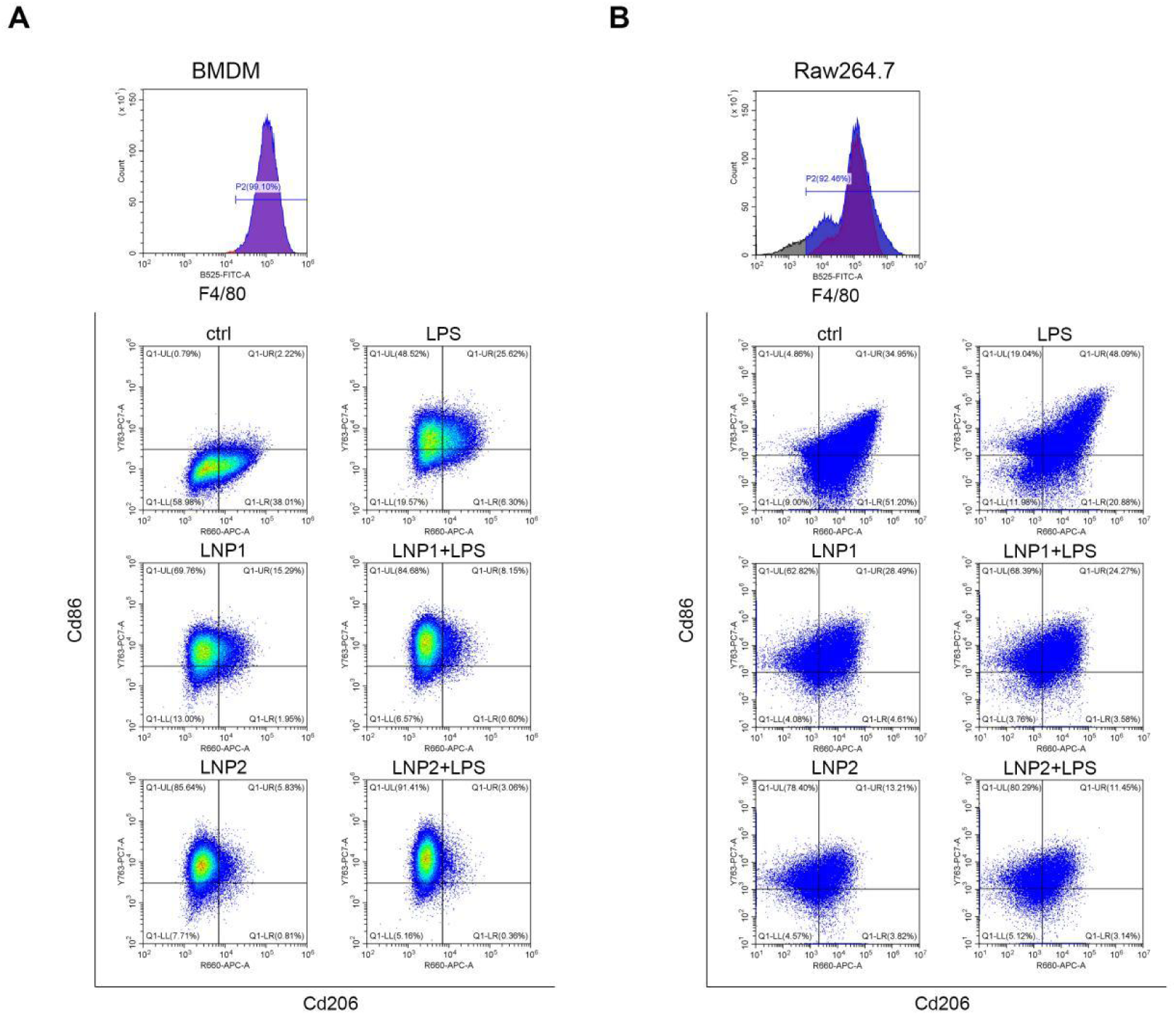
LNP1/2 transfection or LPS treatment promotes M1 polarization of macrophages. (A, B) Flow cytometric analysis of Cd206 and Cd86 expression in BMDM or Raw264.7 cells following treatment with LNP or LPS. Cd86/Cd206-positive cells were selected from the F4/80^+^ population.

**Figure S4.**
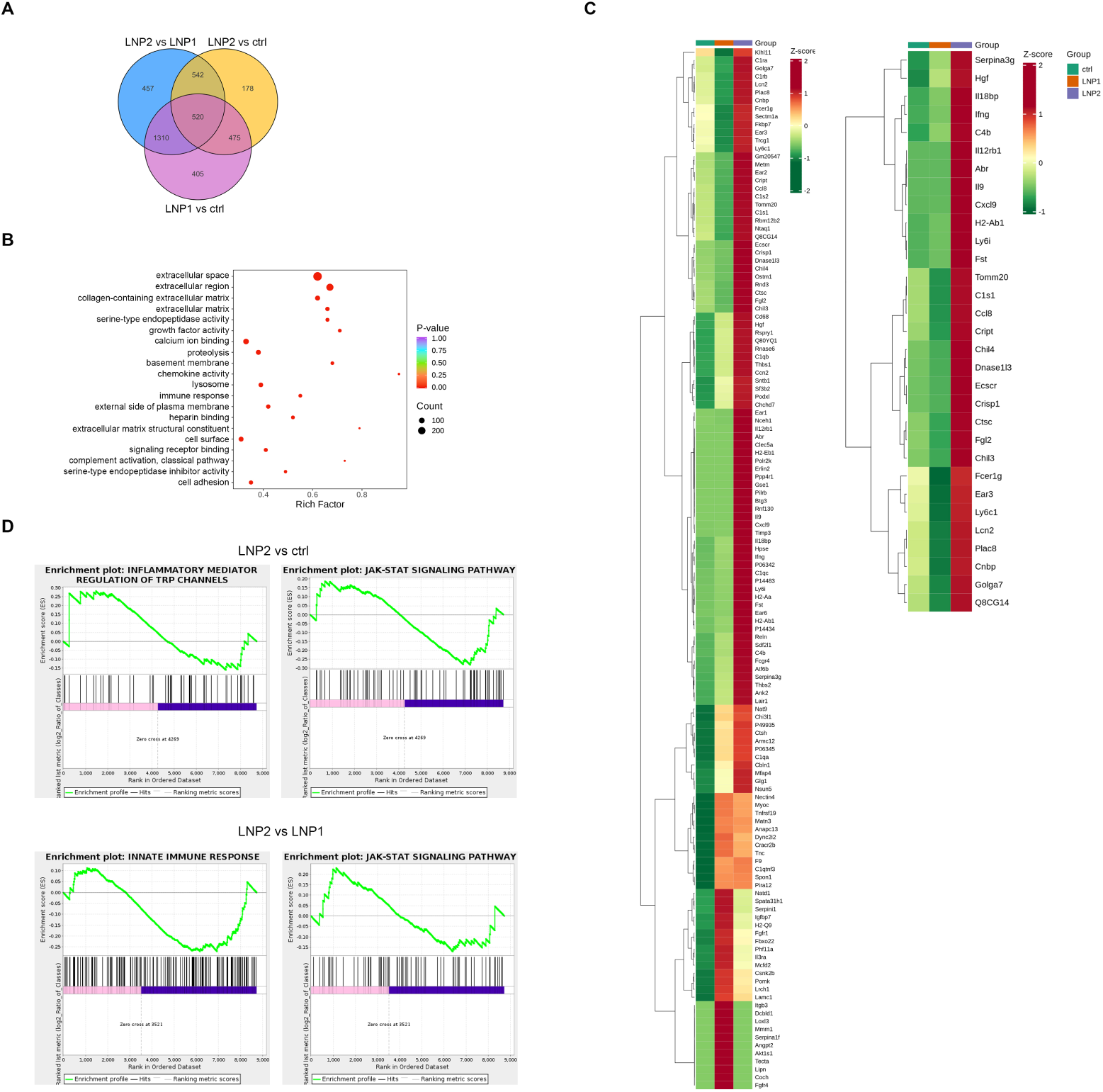
Mass spectrometry profiling of secretory proteins from BMDM cells armed with LNP1 or LNP2. (A) Venn diagram illustrating the overlap and differences in mass spectrometry results across groups. (B) Gene ontology (GO) analysis of DEPs among groups. (C) Heatmap of DEPs showing at least two-fold change between the LNP2 treatment group and other groups. The left panel highlights the top enriched secretory proteins in LNP2-armed CAR-M compared to the control group, while the right panel shows those highly enriched in LNP2-armed CAR-M relative to LNP1-armed CAR-M. (D) GSEA analysis of DEPs comparing LNP2 with control or LNP2 with LNP1 treatment groups.

**Figure S5.**
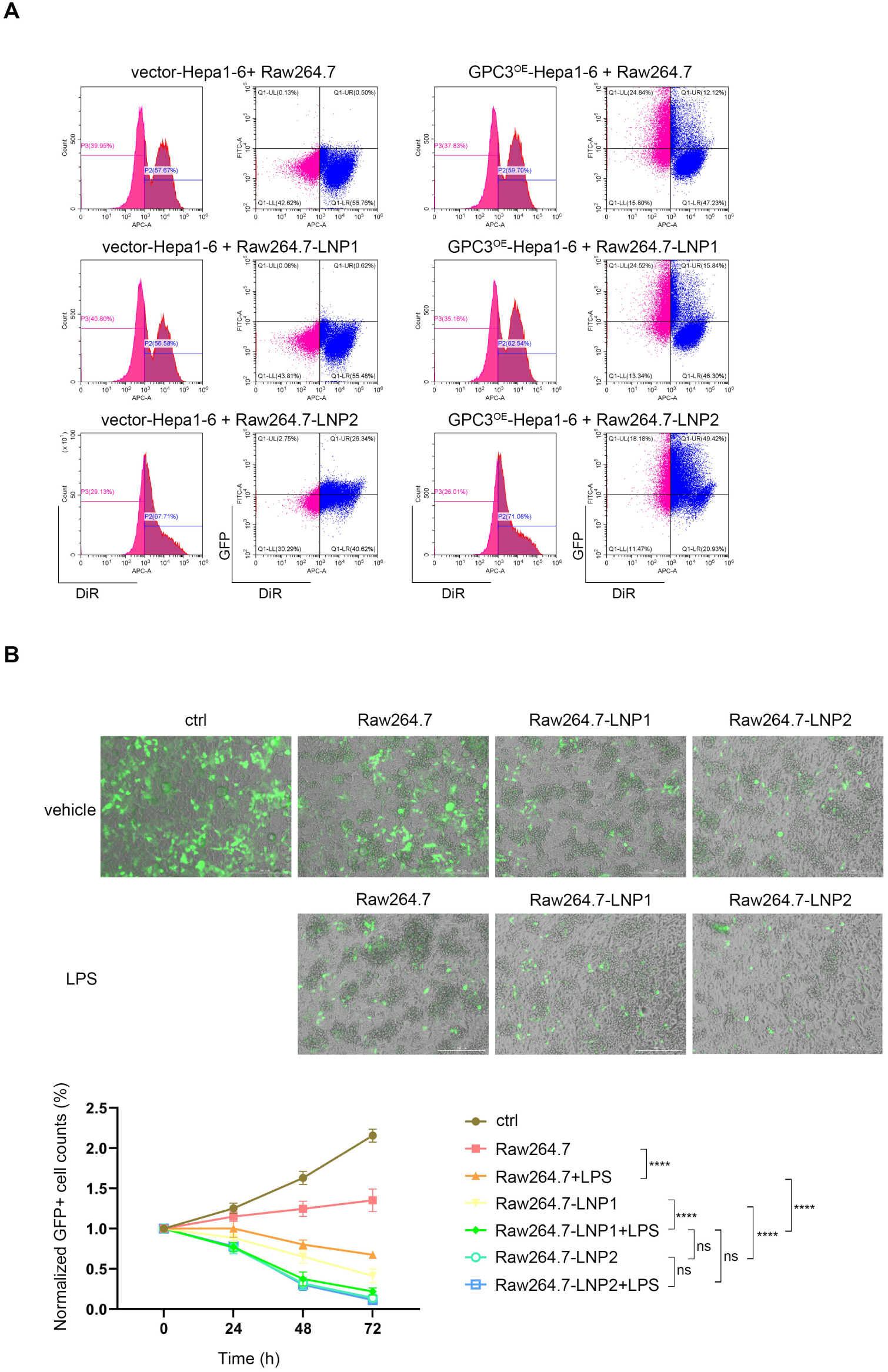
CAR-M cells demonstrate potent anti-tumor activity in vitro. (A) Flow cytometric results showing DiR^+^ or GFP^+^ cells after 6 hours of co-culture between Hepa1-6 and Raw264.7 cells. The left panel displays results from control Hepa1-6 cells transfected with a GFP vector, while the right panel shows results from GPC3-2A-GFP-overexpressing Hepa1-6 cells. (B) Live-cell imaging of GPC3^OE^-Hepa1-6 cells after 72 hours of co-culture with CAR-M cells at a 1:1 E:T ratio, along with time-course quantification of GFP^+^ cells across treatment groups. The statistical data were presented as mean ± s.e.m., and two-way ANOVA analysis was used to calculate the significance. \*\*\*\**p* < 0.0001. ns, not significant.

**Figure S6.**
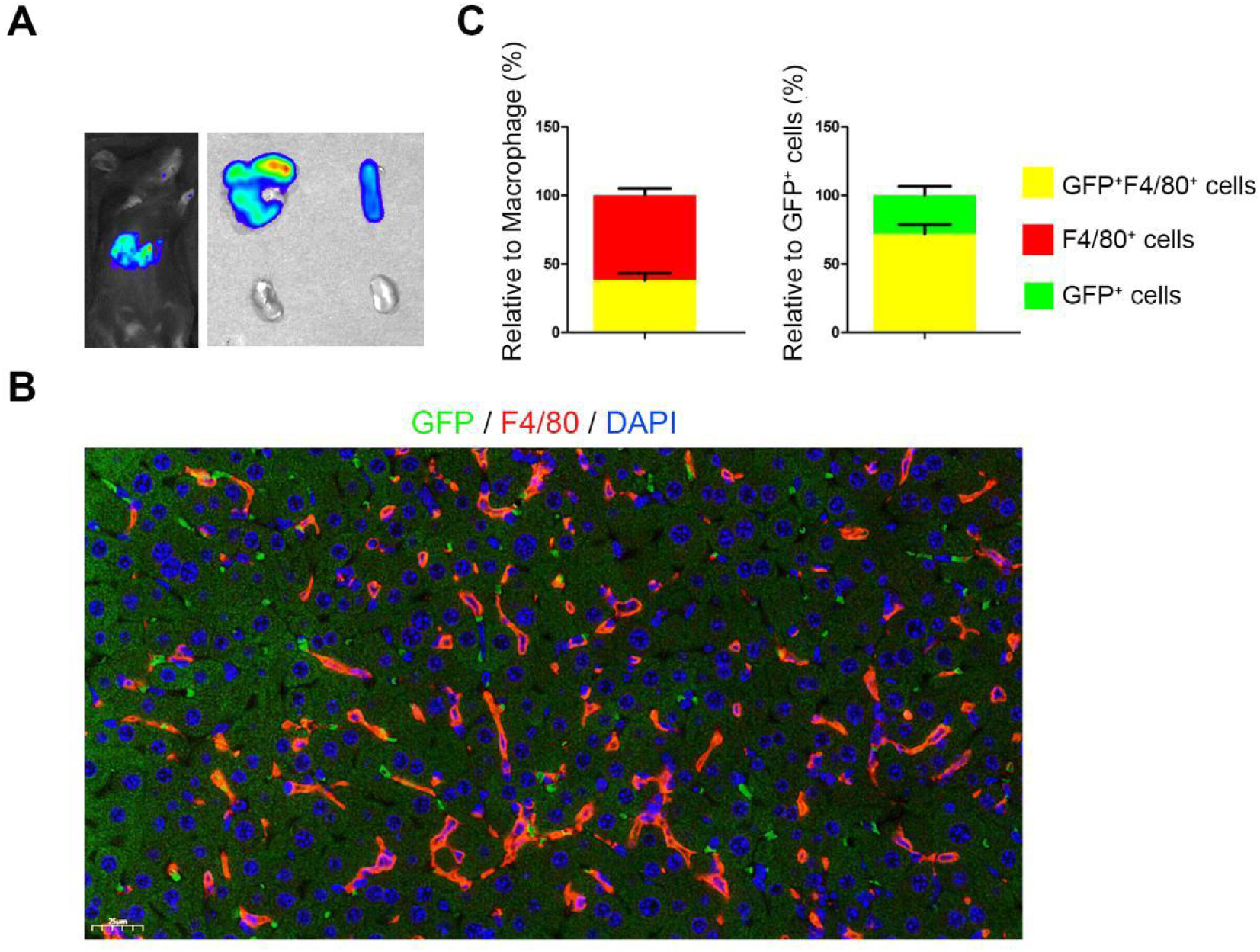
Intravenous injection of LNP targets the liver in mice in vivo. (A) Bioluminescence signal of mice 4 hours after intravenous injection of LNP-luciferase. The dosage of LNP-mRNA was approximately 5 μg diluted in 200 μL of PBS solution. The right panel shows the distribution of the signal across different organs, including the liver, spleen, and kidney. (B) Immunofluorescence image of GFP signals in mice liver tissues after intravenous injection of LNP-eGFP. The liver tissues were dissected, fixed, paraffin-embedded, sectioned, and then subjected to immunofluorescence labeling of F4/80 cells. The spatial relationship between GFP-expressing cells and Kupffer cells/macrophages was subsequently analyzed. (C) Stacked bar chart showing the proportion of GFP^+^ F4/80^+^ cells among total GFP^+^ cells or total F4/80^+^ cells. Fluorescence signals were quantified using ImageJ software.

**Figure S7.**
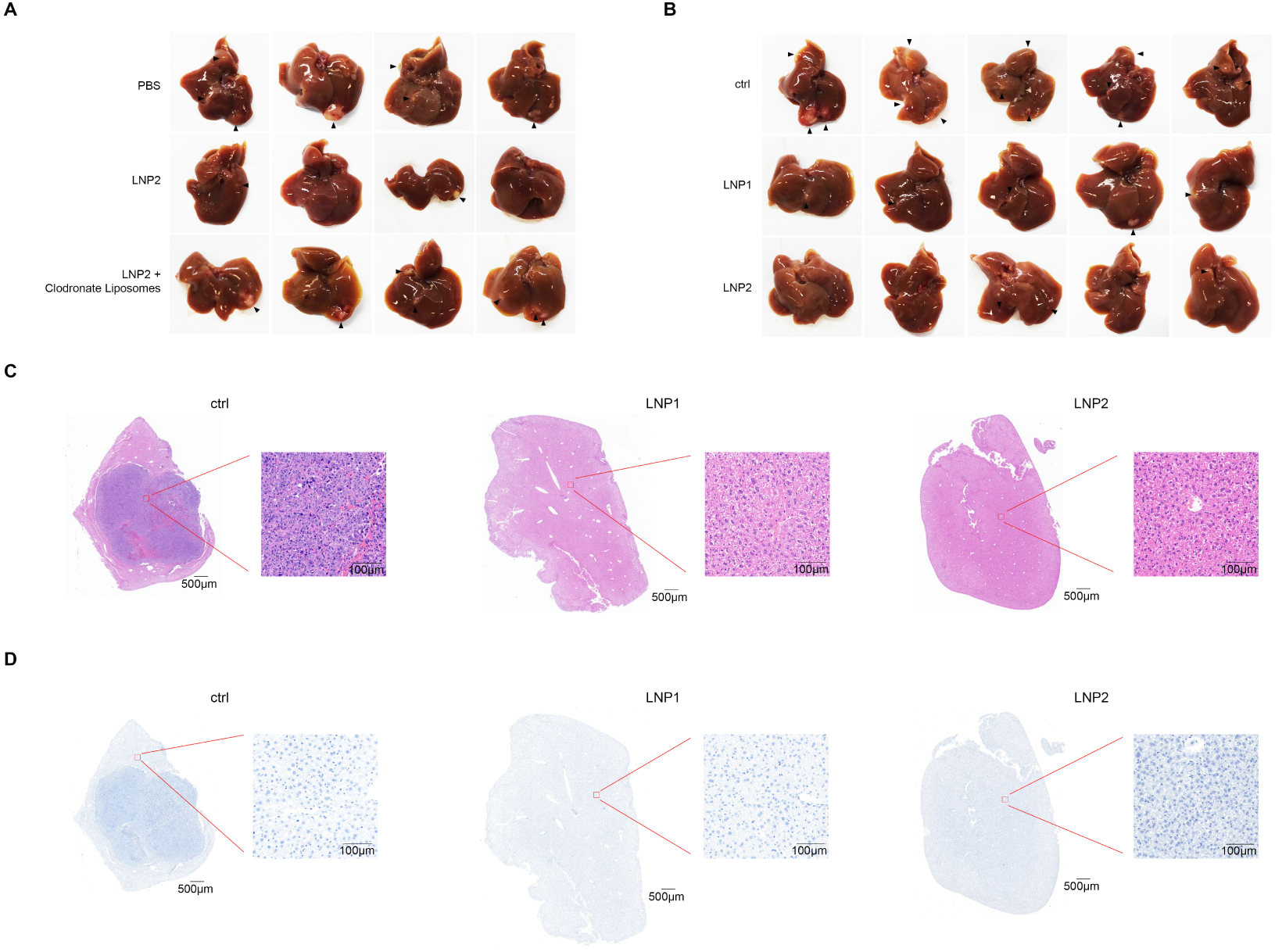
CAR-Ms generated by in vivo LNP delivery suppress tumor growth in the mouse liver. (A, B) Dissected liver organs from orthotopic xenograft tumor-bearing mice after administration of LNP1/2 or clodronate liposomes. Black arrows indicate tumor lesions in the liver. (C, D) Representative images of H&E and p-Stat1 immunohistochemical staining from liver tissues of the aforementioned mouse model are shown. The rectangle indicates a locally magnified area. In the control group, both the tumor region and its surrounding microenvironment were visible within the same field of view. p-Stat1 expression was specifically detected in the tumor microenvironment.

**Figure S8.**
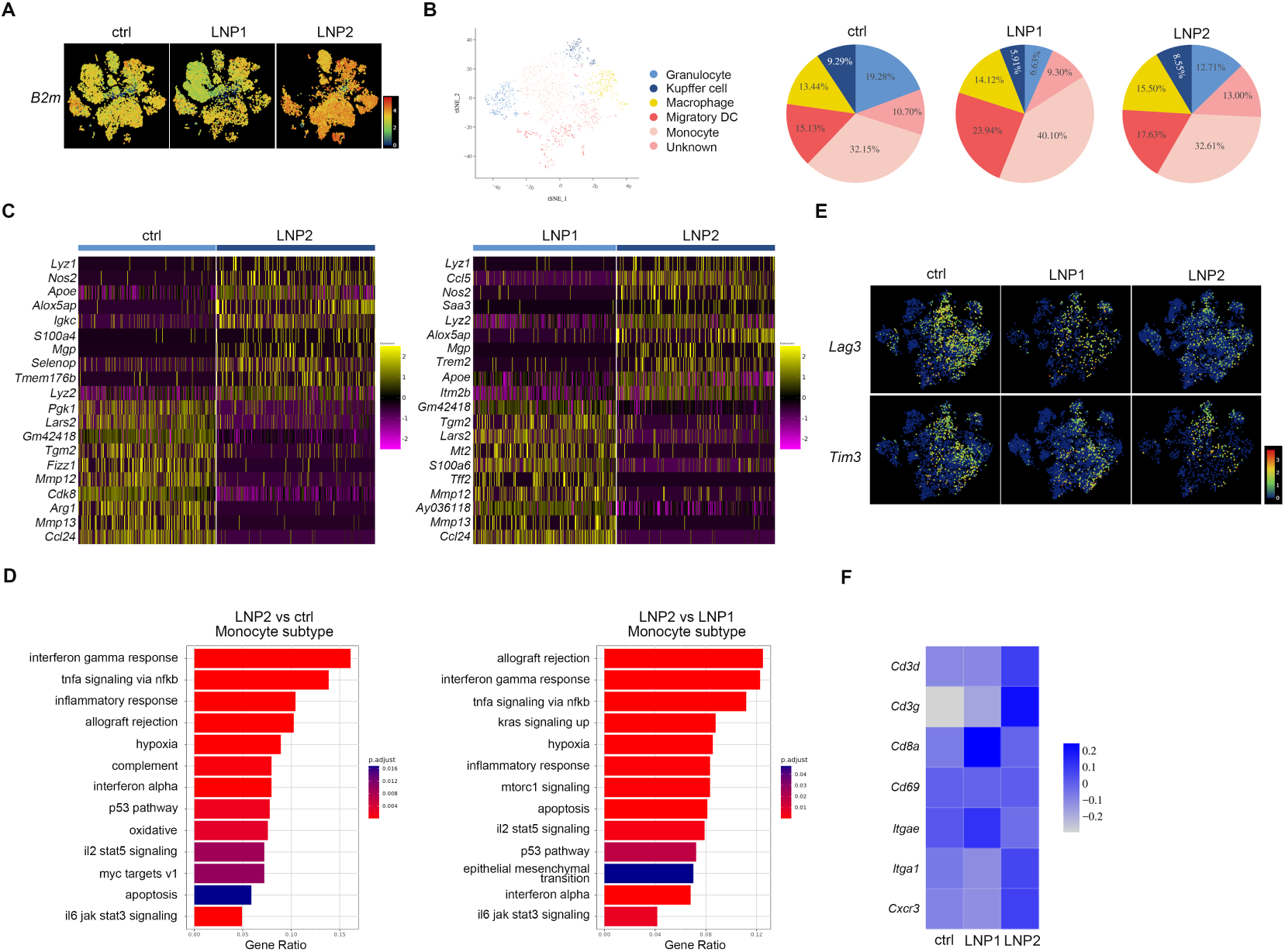
LNP2-armed CAR-Ms remodel the immune microenvironment into a pro-inflammatory and anti-tumor state. (A) mRNA expression levels of *B2m* in the TME across different samples, with color intensity representing gene expression levels. (B) t-SNE plot and pie chart depicting the distribution of cell subsets within the monocyte group across different samples. (C) Heatmap analysis showing the most enriched genes in the monocyte group from LNP2-treated mice compared to control or LNP1-treated mice. (D) Bar plot displaying the most significantly enriched Hallmark gene sets differentially expressed in the monocyte group between LNP2 and control or LNP2 and LNP1-treated mice. Hallmark gene sets were obtained from the MSigDB database. (E) mRNA expression levels of *Lag3* and *Tim3* in the TME across different samples, with color intensity reflecting gene expression levels. (F) Heatmap analysis of the expression of effector memory T cell-related genes across different samples.

**Figure S9.**
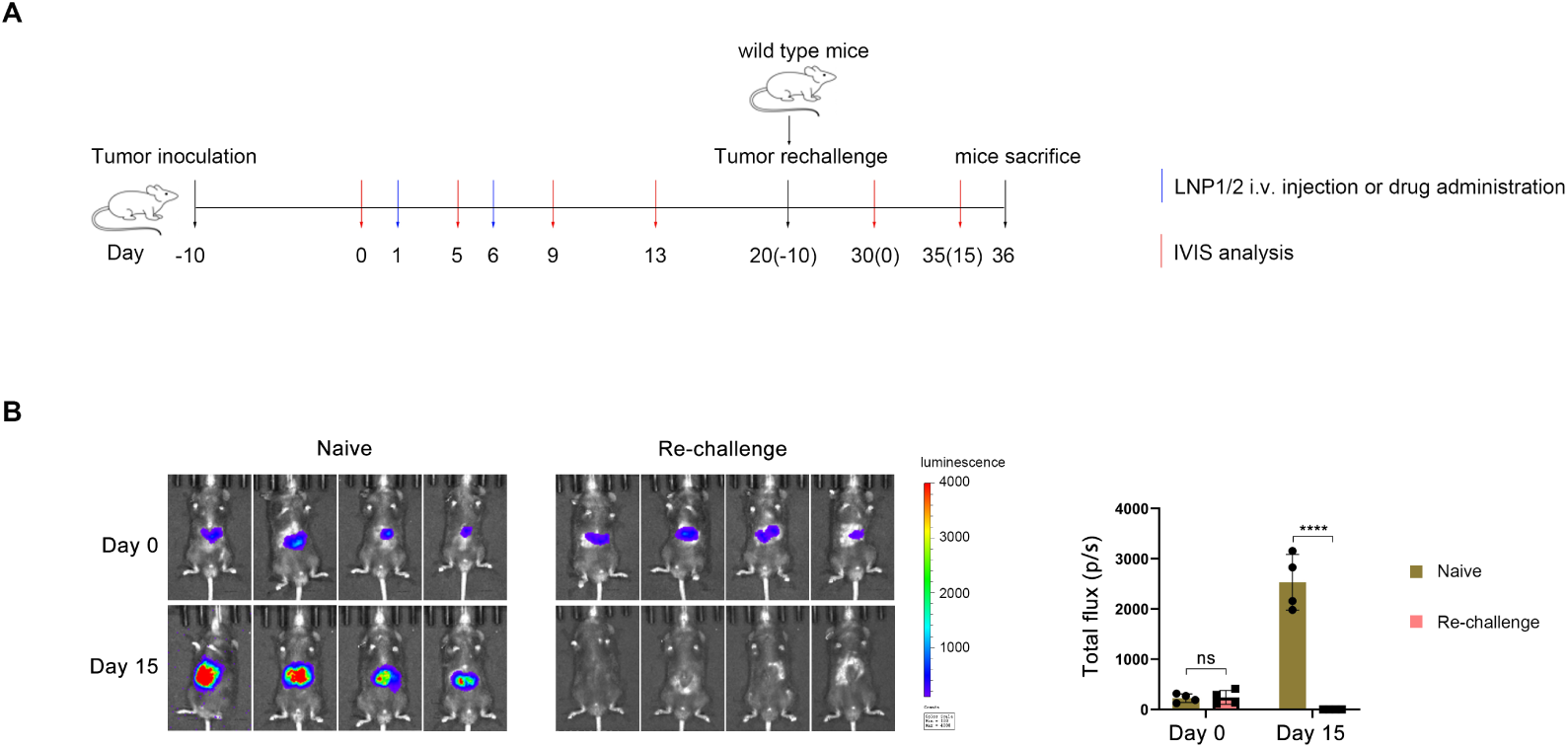
LNP2 pre-treatment enables mice to establish immune memory against GPC3-overexpressing tumor cells. (A) Schematic illustration of the experimental design for the tumor re-challenge assay in a liver orthotopic xenograft tumor model. (B) In vivo bioluminescence imaging showing time-dependent tumor growth in mice across different treatment groups. The right panel presents the quantification of bioluminescence signals across treatment groups.

**Figure S10.**
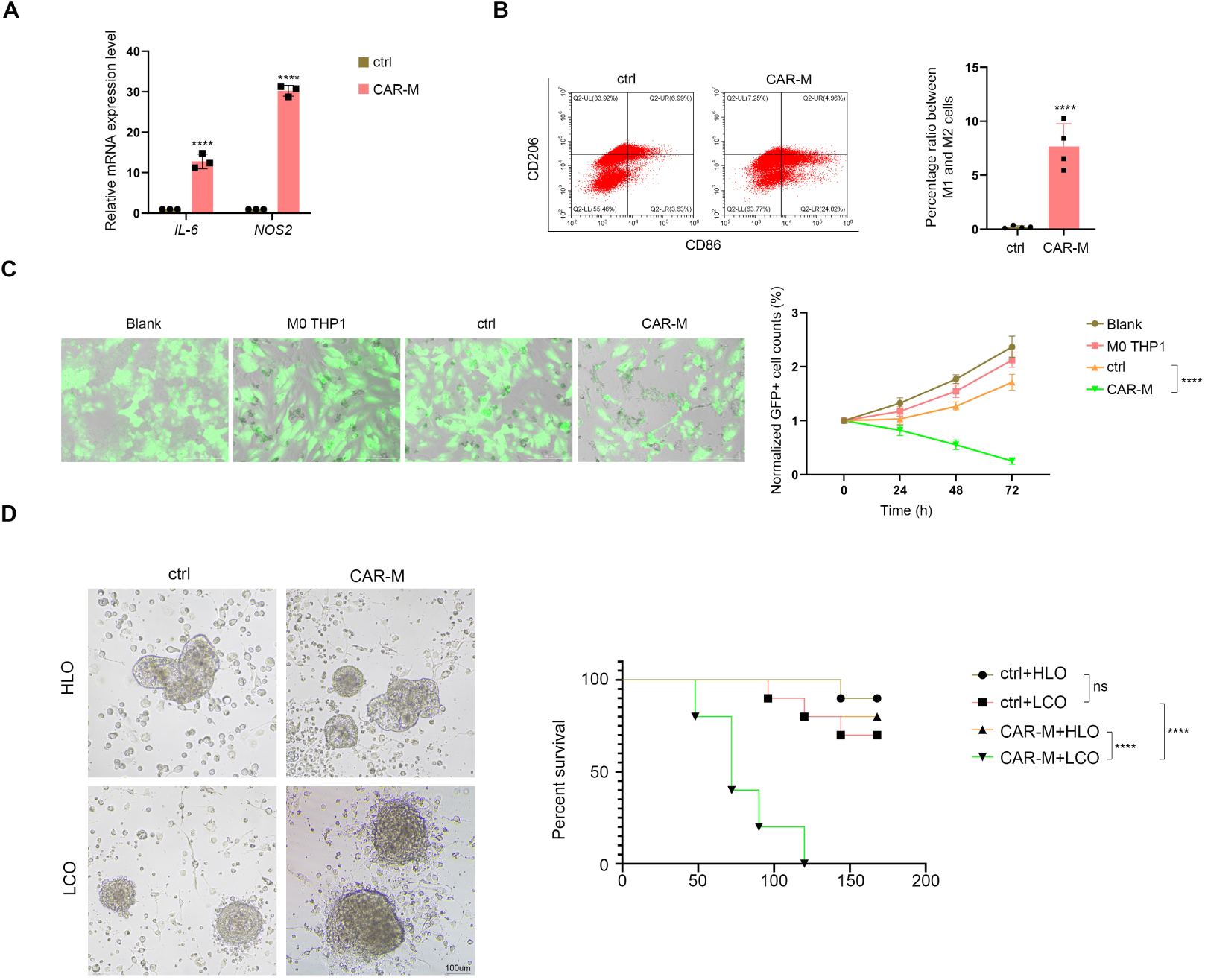
CAR-Ms produced by LNP transfection efficiently eliminate human liver cancer cells in vitro. (A) qRT-PCR detection of M1 marker expression in PBMC-derived macrophages (ctrl) or CAR-Ms. (B) Flow cytometric evaluation of CD86 and CD206 expression in control macrophages or CAR-Ms, along with the corresponding quantification of M1/M2 ratios. (C) Live-cell imaging of GFP-overexpressing Huh7 cells after 72 hours of co-culture with THP1 or CAR-M cells. No THP1 cells were added in the blank group. M0 THP1 cells refer to THP1 cells without activation factor treatment, whereas the ctrl group refers to THP1 cells pre-treated with LPS and IFN-γ. The right panel shows the time-course quantification of GFP^+^ cells across treatment groups. (D) Morphological dynamics of HLO or LCO co-cultured with control or THP1-derived CAR-M cells over time. The organoids shown are representative of at least 30 organoids per group. The right panel presents Kaplan-Meier survival analysis of HLO or LCO after long-term co-culture with THP1 control or CAR-M cells. The co-culture system was observed and recorded every 24 hours. Organoids exhibiting apparent disaggregation or disruption were considered as death events. Statistical data were presented as mean ± s.e.m., and t-tests analysis or two-way ANOVA analysis was used to calculate the significance. \*\*\*\**p* < 0.0001. ns, not significant.

