## Supplementary figures and images for "An in situ generated CAR-M with IFN-γ and negative dominance Sirpα isoform augments hepatocellular carcinoma immunotherapy"

### Supplementary Figures

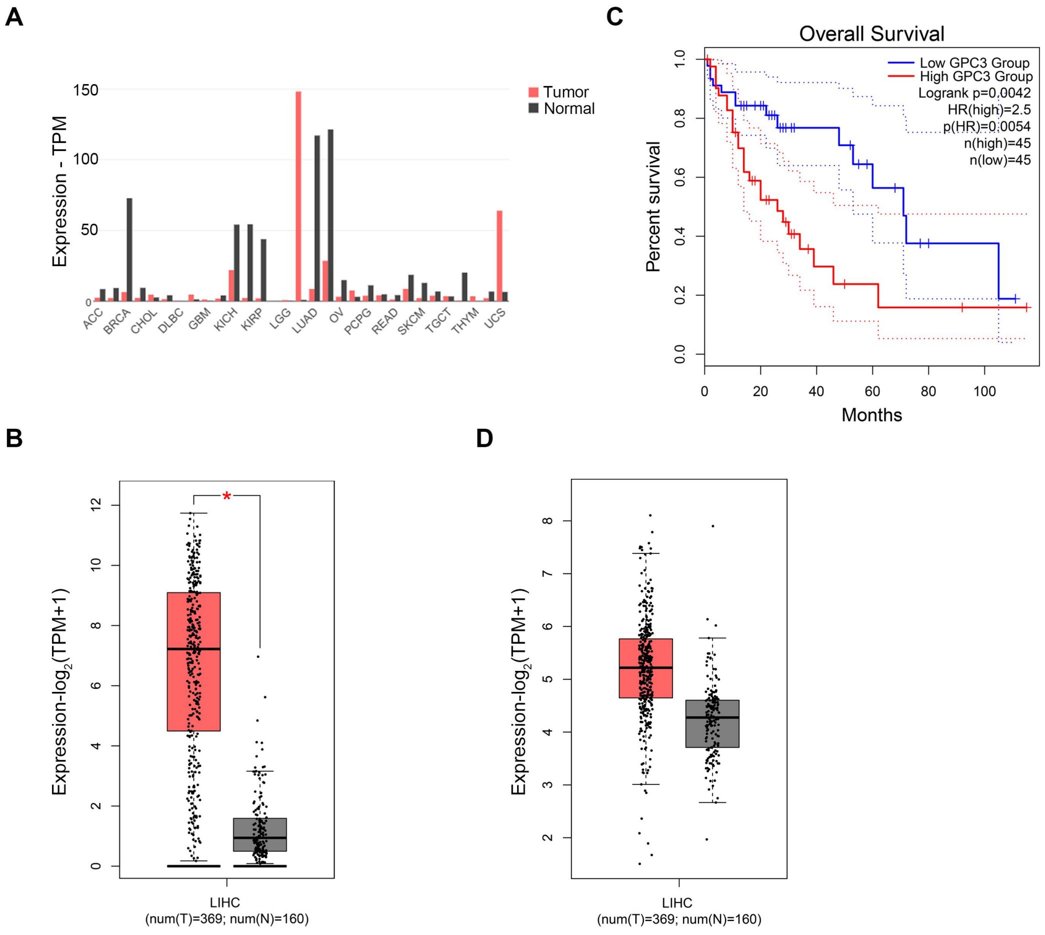

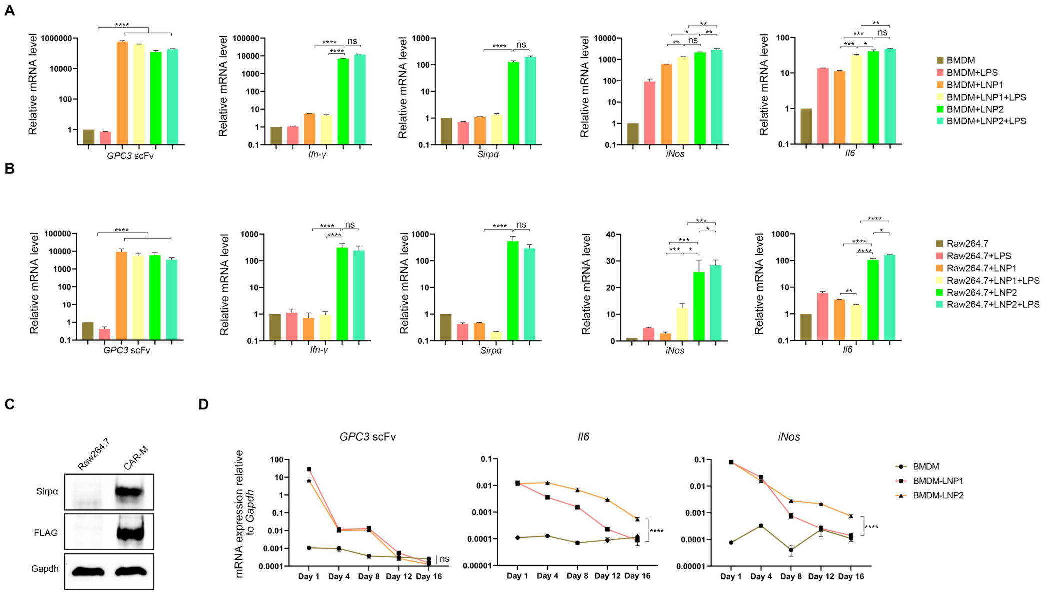

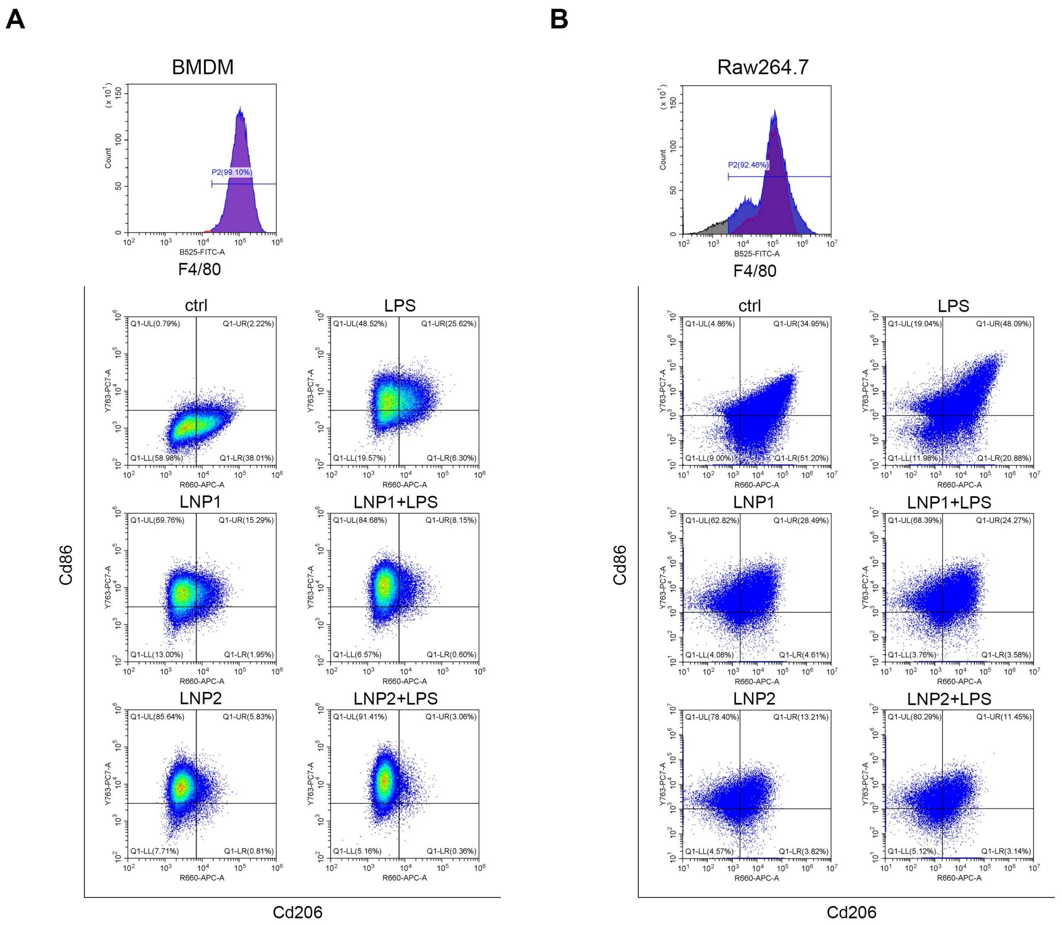

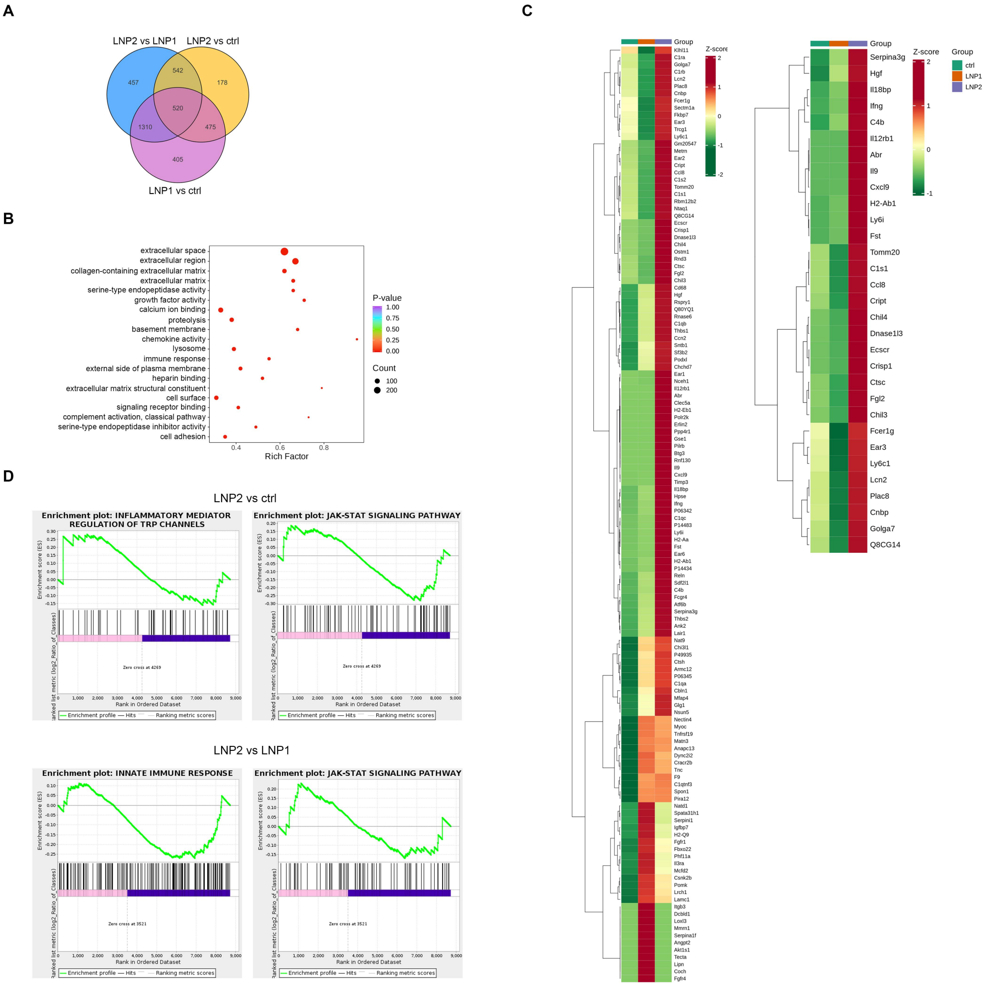

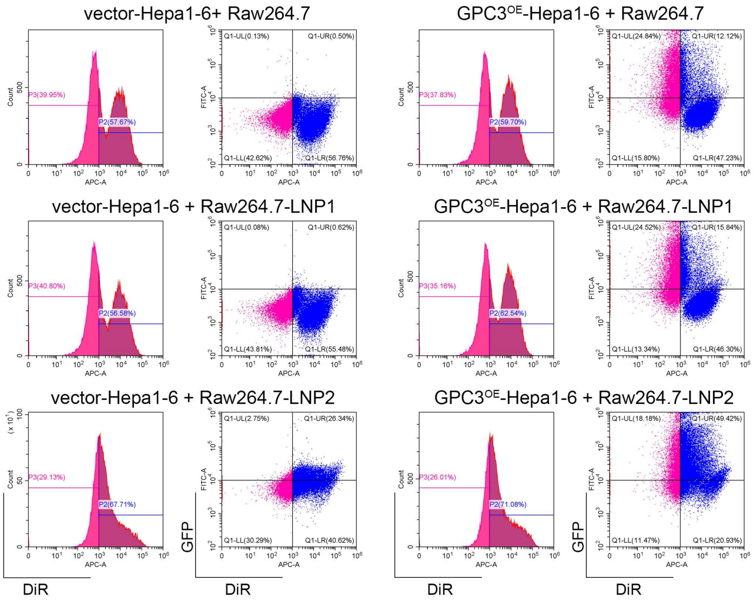

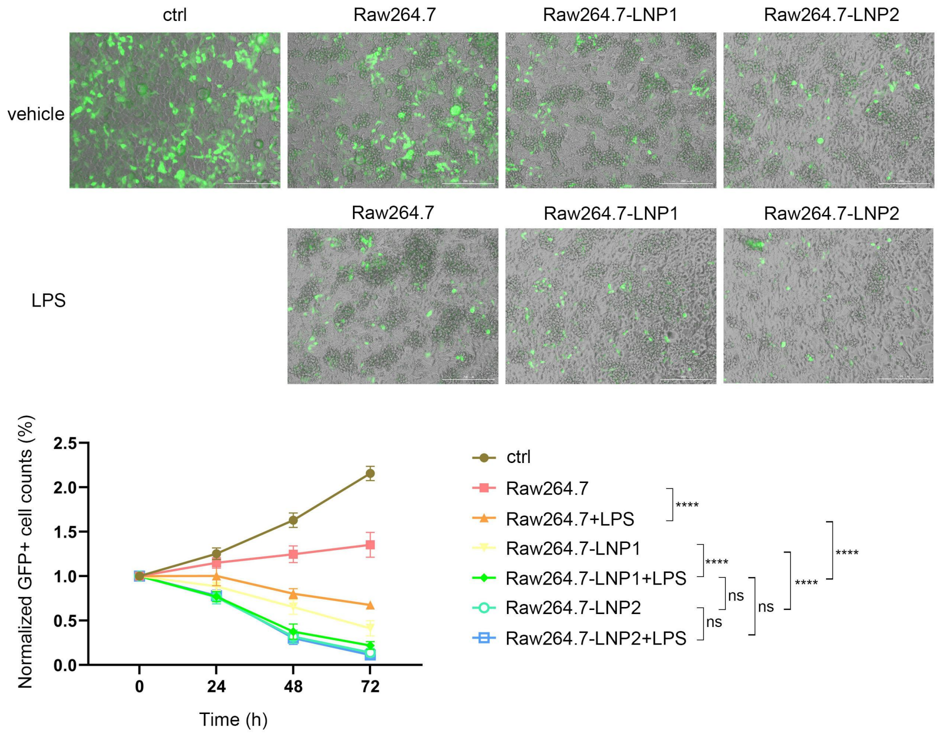

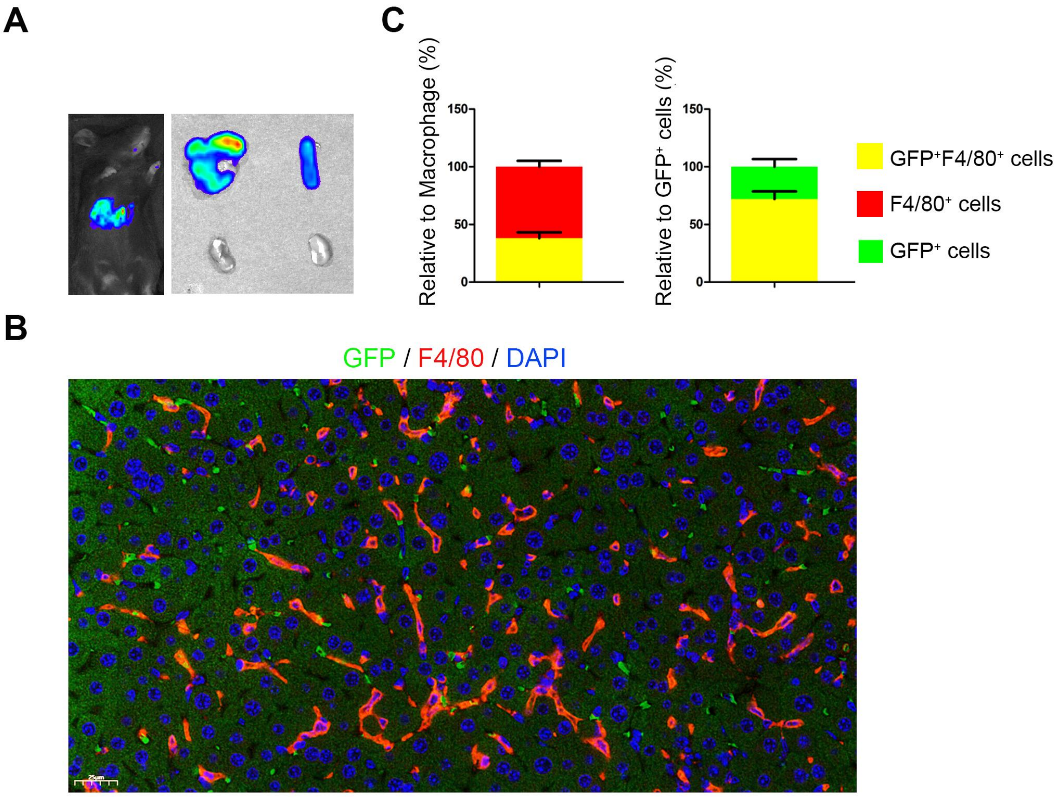

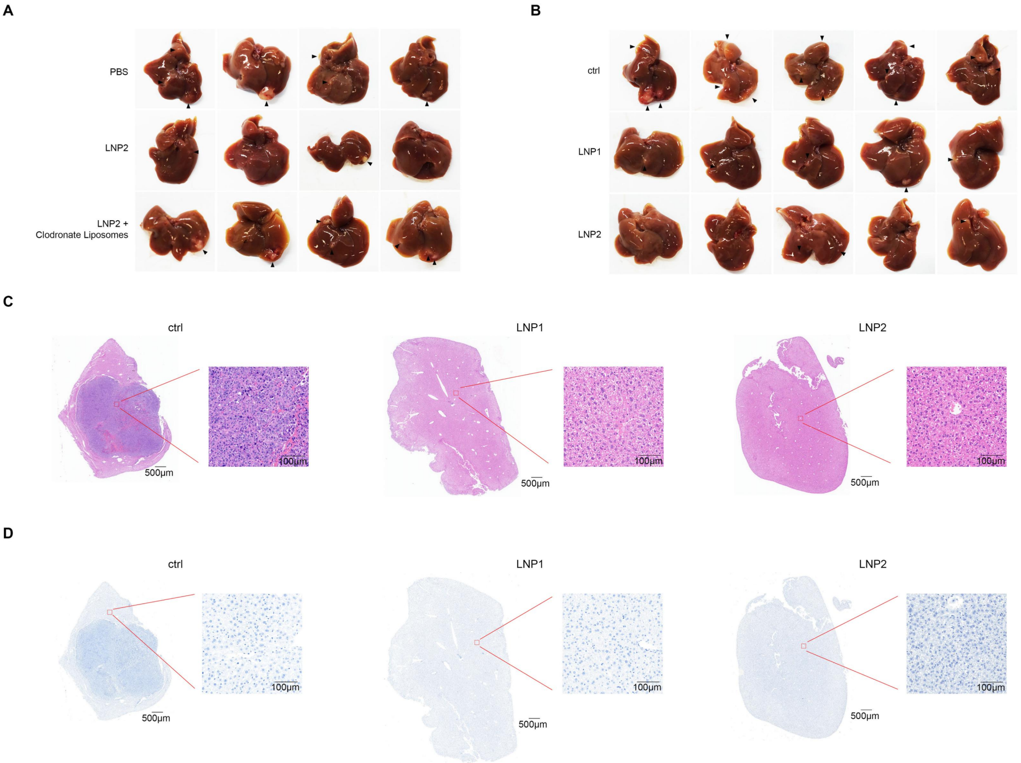

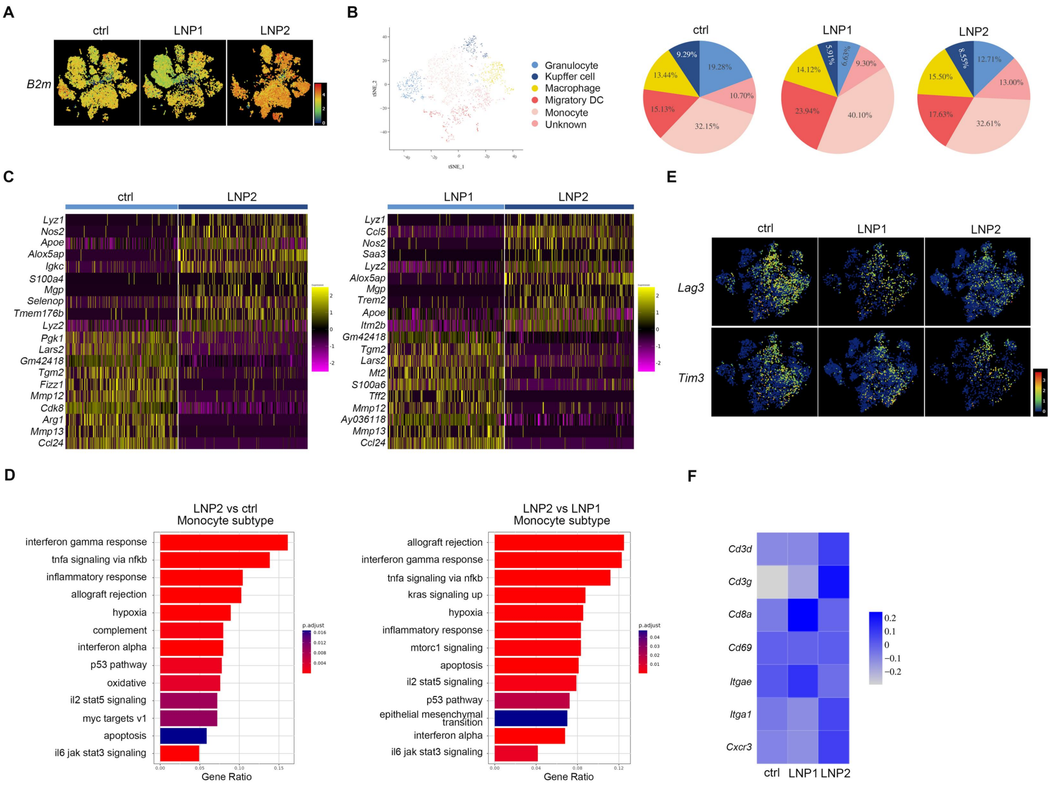

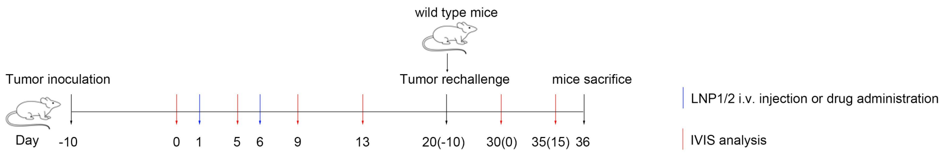

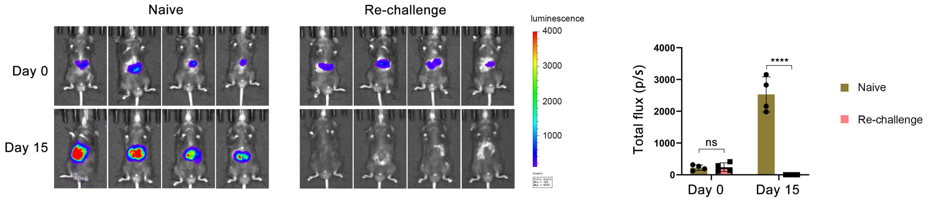

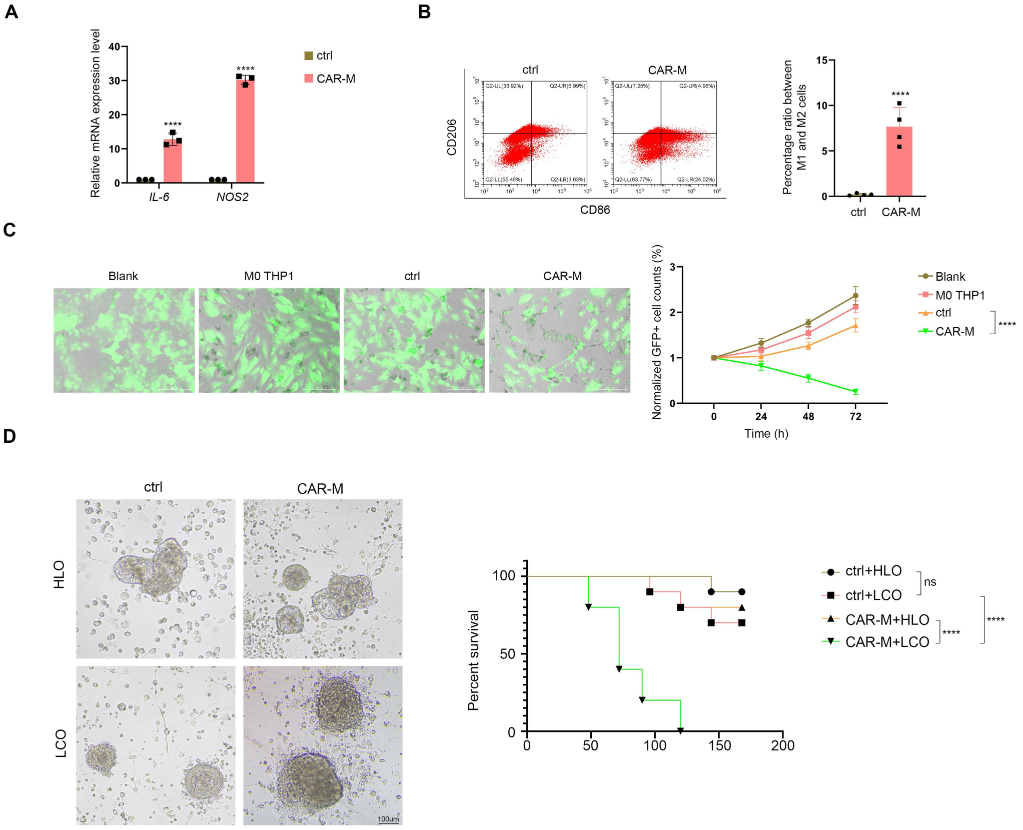
